# Axonopathy and altered synaptic development in early hippocampal epileptogenesis of Dravet syndrome

**DOI:** 10.1101/2023.10.04.560735

**Authors:** Nikolas Layer, Peter Müller, Maya Ayash, Friederike Pfeiffer, Meret Saile, Fabian Klopfer, Stefano Iavarone, Andrea Santuy, Petra Fallier-Becker, Ulrike B. S. Hedrich, Holger Lerche, Henner Koch, Thomas V. Wuttke

## Abstract

Dravet syndrome caused by *SCN1A* variants is a severe developmental epileptic encephalopathy (DEE) characterized by pharmaco-resistant epileptic seizures and progressive neurodevelopmental decline with cognitive impairment and autism-spectrum-traits. Numerous preceding studies indicate that the initial pathophysiology due to impaired Na_V_1.1 function mainly derives from reduced interneuron firing leading to a network hyperexcitability (Bender et al. 2012). However, little is known how epileptogenesis and generally disease pathogenesis progress from the inborn molecular defect to infantile seizure onset. We address this question in a Dravet mouse model by comprehensive single-cell RNA sequencing and selected downstream analysis via single-cell electrophysiology, histology, live cell imaging and electron microscopy. Our data reveal a continuum of early primary (preseizure) and secondary (post-seizure onset) transcriptomic changes in various cell populations in the hippocampus. Focusing on *cornu ammonis*, we find a number of transcriptional pathways that are dysregulated including synaptic transmembrane adhesion molecules of the neurexin superfamily and voltage-gated ion channels. Further investigations support an ultrastructural and functional axonopathy and synaptopathy of parvalbumin interneurons. These processes precede somatic firing impairment and seizures suggesting they underlie fundamental early-phase disease mechanisms. Taken together we provide a cellularly resolved transcriptomic resource of early disease phases of Dravet syndrome and demonstrate epileptogenesis beyond Na_V_1.1 loss-of-function during an early developmental time window of CNS maturation. Altogether these data establish proof-of principle that the concept of epileptogenesis, originally devised for acquired forms of epilepsy, similarly applies to genetic epilepsies and DEEs.

## Introduction

Dravet syndrome (DS) is a therapy-resistant developmental epileptic encephalopathy characterized by intractable seizures, cognitive impairment, autistic features, motor symptoms and increased mortality with a symptom onset earlier than one year of age. Seizure types in DS range from febrile and afebrile, focal clonic, and generalized clonic seizures to myoclonic and atypical absence seizures (Zuberi et al. 2022). The majority of cases are caused by heterozygous *de novo* loss-of-function (LOF) variants in *SCN1A*, encoding for the voltage-gated sodium channel Na_V_1.1 (Claes et al. 2001; Dravet und Oguni 2013). Genetic Dravet mouse models harboring variants in *Scn1a*, which either lead to haploinsufficiency by reduced protein expression or altered channel gating, closely recapitulate the typical clinical presentation and disease progression from a presymptomatic to a worsening/severe disease phase which finally transitions into the stabilization phase. Whereas steadily increasing Na_V_1.1 expression levels from the second week of age (Cheah et al. 2013) on are only reflected phenotypically in a higher susceptibility to febrile seizures during the preseizure phase in a LOF *Scn1a* mouse model (Dutton et al. 2017), the high frequency seizure phase is characterized by spontaneous seizures and emerging comorbidities like motor dysfunction, impaired social behavior and an increased risk of seizure-associated sudden death (Ricobaraza et al. 2019; Fadila et al. 2020; Bahceci et al. 2020; Gerbatin et al. 2022). Characteristically, first convulsions in patients are febrile seizures which are thought to emerge from hippocampal areas implying initial hippocampal disbalance of excitation and inhibition (E/I) (Patterson et al. 2017; Kloc et al. 2023; Dutton et al. 2017; Hesdorffer et al. 2013). In this context, multiple electrophysiological studies of the hippocampus demonstrated that Na_V_1.1 dysfunction in fact mainly affects GABAergic interneurons (Yu et al. 2006; Almog et al. 2021; Stein et al. 2019). Amongst these, parvalbumin fast-spiking interneurons (fsIN) are known to be especially vulnerable as reflected by a reduction of somatic action potential firing at high current injections (Hedrich et al. 2014; Mattis et al. 2022). After transitioning into the stabilization phase, both seizure frequency and the rate of sudden death decline, although the comorbidities persist to adulthood (Oakley et al. 2011; Ricobaraza et al. 2019). Whereas the latter symptoms were initially believed to be a pure consequence of recurrent severe seizure activity, more recent studies in several models of epileptic encephalopathy (including DS) found an onset of at least a subset of comorbidities independent of and even before seizures (Fadila et al. 2020). As both seizures and comorbidities emerge between the second and third week in Dravet mice during a time window of ongoing CNS circuit formation, malfunctioning synaptogenesis, altered circuit maturation and pathologic network remodeling could represent underlying molecular disease mechanisms (Mantegazza und Broccoli 2019). However, prior work did not systematically address these hypotheses but remained largely confined to the initial genetically determined ion channel impairment (i.e. loss-of-function of the Na_V_1.1 sodium channel in DS) and the immediate consequences for neuronal properties and function. Along these lines, very recent studies explored the therapeutic potential of *post hoc* restoration of Na_V_1.1 function via up-regulation of *Scn1a* gene expression. Although, this approach achieved amelioration of seizures and mortality (Fadila et al. 2023; Han et al. 2020; Valassina et al. 2022) it failed to revert the neurodevelopmental phenotype in its entirety, thereby suggesting additional pathophysiological mechanisms beyond the ion channel defect.

How initial channel and interneuron dysfunction in DS may interfere with developmental transcriptional processes in interneurons and other CNS cell types is currently unknown. However, it is conceivable that perturbations at this level may set off or act as early (primary) epileptogenic mechanisms which in turn may trigger a cascade of secondary pathogenic changes altogether leading to the developmental and epileptic encephalopathy phenotype of DS. As next generation sequencing technology has majorly advanced over the last years, research is now able for the first time to study disease pathogenesis and specifically epileptogenesis beyond the initial molecular hit in thousands of cells in parallel on the transcriptomic level with cellular resolution. Here, we focused our study on a mouse model of DS harboring the human recurrent missense variant A1783V (Depienne et al. 2009; Klassen et al. 2014). This variant alters the voltage-dependence of activation and slow inactivation, without affecting peak sodium current density (Layer et al. 2021). We performed single nuclei RNA-sequencing (snRNAseq) in the hippocampus of Dravet animals at P16 (just before seizure onset) and at P20 (shortly after seizure onset) as well as in respective control wildtype (wt) animals to explore pathways of primary and secondary hippocampal epileptogenesis and to capture the dynamics of disease mechanisms across virtually all neuronal and non-neuronal cell types. Combining differential gene expression and pseudotime trajectory analysis revealed multiple primary and secondary transcriptomic signatures, particularly in PV interneurons (PVIN), which progressively evolve on a single cell level from an initial healthy to a pathological state. Functional and structural investigations revealed an axonal and synaptic dysfunction of fsIN preceding the previously described somatic firing deficit and identify fsIN axono- and synaptopathy as early/initial hippocampal mechanisms of epileptogenesis in DS.

## Results

### Single nuclei RNA sequencing in *Scn1a*^+/A1783V^ mice reveals changes in neurodevelopmental processes in the hippocampus

Pathophysiological alterations in Dravet mouse models including heterozygous *Scn1a*^+/A1783V^ knock-in mice have mainly been investigated via single cell electrophysiology (Almog et al. 2021; Layer et al. 2021), surface EEG recordings (Fadila et al. 2020; Kuo et al. 2019), histology (Martín-Suárez et al. 2020; Ricobaraza et al. 2019) and bulk proteomics (Miljanovic et al. 2021). However, these approaches are low throughput and provide only limited insight into cellular mechanisms of disease pathogenesis and epileptogenesis or don’t have a sufficient resolution to understand cellular pathomechanisms of Dravet syndrome. To overcome these limitations, we captured a comprehensive data set with single cell resolution which covers early pathophysiological processes in thousands of neuronal and non-neuronal cells by single-nuclei RNA-sequencing. Hippcampal tissue was collected before (P16) and after (P20) seizure onset (Figure 1A) from *Scn1a*^+/A1783V^ mice and compared to age-matched wildtype (wt) littermate controls. After quality control, doublet removal and dataset integration, we first identified all major neuronal and glial cell types based on known marker genes in all biological replicates (Figure 1B, D). Since the resolution of UMAP-clustering based on the nearest neighbor algorithm was not sufficient to yield separation of different GABAergic interneuron subtypes, we performed separate clustering for all nuclei clusters expressing the interneuron marker genes *Gad1* and *Gad2* which allowed identification of parvalbumin (PV), somatostatin (SST), vasoactive intestinal peptide (VIP) expressing and further interneuron types (Figure 1C, E). For PVINs, we found a lower overall number of cells (than expected in relation to the numbers of detected SST and VIP neurons) based on parvalbumin mRNA transcripts at both time points, in line with PV expression being an unreliable marker due to its activity-dependence and relatively late onset of expression only from about two weeks of life on (Lecea et al. 1995; Vormstein-Schneider et al. 2020). A second well-known marker gene for the PV cell population is the K_V_3.1 potassium channel subunit encoding gene *Kcnc1* (Chow et al. 1999) which enabled reliable clustering and identification of PVINs in our datasets. Within the neuronal populations we found no change in the expression of *Scn1a* between both genotypes at both timepoints (data not shown) consistent with the missense type of the p.(Ala1783Val) variant and with previously published Western blot data (Ricobaraza et al. 2019).

**Figure 1.**
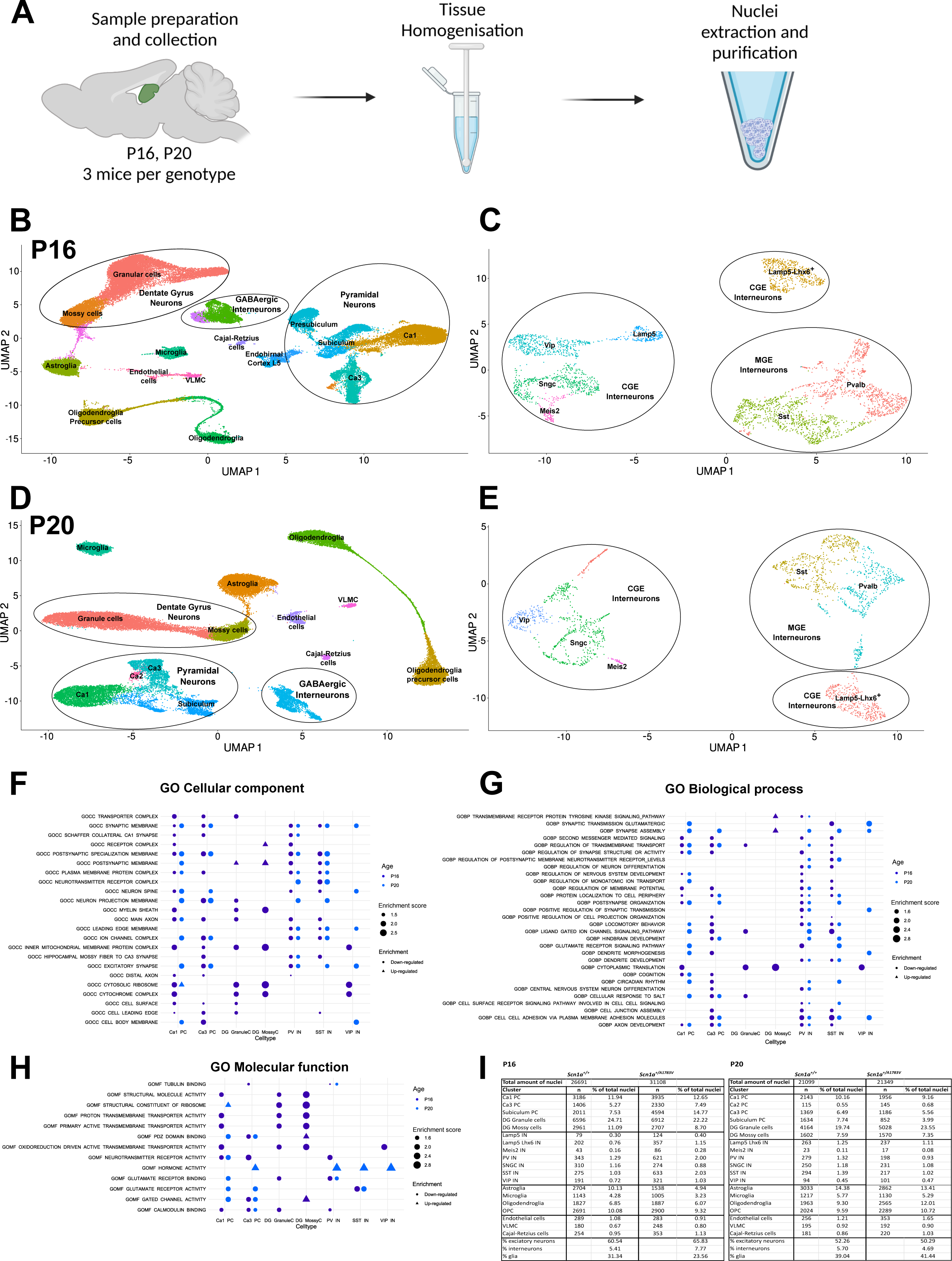
Single nuclei RNA-Sequencing in *Scn1a*^+/A1783V^ mice before and after seizure onset. **A:** Schematic representation of workflow for single nuclei RNA (snRNA)-sequencing in hippocampal tissue. **B, D**: Umap representation of P16 and P20 hippocampal snRNA data in *Scn1a*^+/+^ and *Scn1a*^+/A1783V^ mice samples with identified cell cluster annotation. **C, E:** Umap representation of P16 and P20 hippocampal snRNA interneuron data subclusters. **F - G**: Dot plots of GO terms (GO cellular component, GO biological process, GO molecular function) enriched in at least two cell types after GSEA in *Scn1a*^+/A1783V^ mice at P16 (navy blue) and P20 (light blue); triangles represent significantly up-regulated GO terms while circles represent significantly down-regulated GO terms. **I**: Number of nuclei per cluster in each dataset split by genotype.

To describe the cellular components and processes, which are directly or indirectly affected by the p.(Ala1783Val) variant, or associated with pathologically altered hippocampal development, we performed Gene Set Enrichment Analysis (GSEA) applying DeSeq2-based testing to all three GO-term domains (Molecular Function, Cellular Component and Biological Process). Next, enriched pathways were extracted based on differentially up- and downregulated genes between age-matched wt and *Scn1a*^+/A1783V^ animals (Figure 1F-H). For both P16 and P20 we found significant pathway enrichment, mostly for downregulated genes. Notably, the number of enriched terms was smaller for dentate gyrus granule and mossy cells than for pyramidal cells (PC) in *cornu ammonis* (CA) 1 and CA3. For PV and SST interneurons, enriched terms were mostly overlapping between cell populations and were already present in the P16 dataset, whereas for CA1 and CA3 PCs we found enrichment of varying pathways at the different timepoints. Most enriched terms in neuronal populations were either connected to the morphologic development of neurons or to synaptic activity and structure. Despite DS having been classically associated predominantly with interneuron dysfunction, we also found pathway alterations in glial populations pointing to more global effects of the disease on the CNS. Noteworthy, there was no upregulation of pathways indicating early neuroinflammation or astrogliosis in glial cells (Supplementary Figure 1 E-G), as the latter had been described for late phases of the pathological development in *Scn1a*^+/A1783V^mice (Martín-Suárez et al. 2020). Furthermore, we neither found relevant differences of the number of nuclei between genotypes at P16 nor at P20 (Figure 1I): Numbers of inhibitory neurons as well as glia in *Scn1a*^+/A1783V^ mice were similar to wt mice at P20, suggesting no increased level of neuronal apoptosis nor increased levels of astrogliosis.

We thus generated a single cell-resolved transcriptomic data set for Dravet syndrome capturing early disease pathogenesis from the preseizure to the high-frequency seizure phase. This data set enables comprehensive mechanistic dissection of DS-associated DEE and will be leveraged throughout this study to uncover mechanisms of epileptogenesis.

### Mild somatic fsIN dysfunction is only present as a transient phenomenon during the high-seizure phase in *Scn1a*^+/A1783V^ mice insufficient to promote pyramidal network hyperexcitability

As GSEA analysis predominantly indicated alterations of pathways associated with excitability of neuronal cell types localized in CA1 and CA3, we next performed somatic whole-cell patch clamp recordings in hippocampal CA1 before seizure onset (P13-15), during the phase of highest seizure frequency (P20-23) and at the stabilization phase (P37-P40). fsIN, known to be especially vulnerable in DS, located at the border between CA1 *stratum oriens* and *stratum pyramidale*, showed similar action potential (AP) properties as expected from previous studies (Goldberg et al. 2010) with a general increase of maximal firing frequencies from the second to the third week of life in wt mice as well as a decrease of input resistance (R_i_), accelerated AP rise times and reduced AP halfwidth (Figure 2F, G, Supplementary statistic table). Whereas passive cell parameters including resting membrane potential (RMP), R_i_ and rheobase were unaltered between fsIN recorded in slices of *Scn1a*^+/A1783V^ and wt mice at all measured time points, the maximum AP frequency of fsIN in *Scn1a*^+/A1783V^ mice was mildly reduced at P20-23 (287.8 ± 16.21 Hz, n = 12 vs. 216.6 ± 18.63 Hz, n = 13, p = 0.0047), yet unaffected at the preseizure phase (Figure 2B, C, Supplementary statistic table). This reduction in AP frequency only became apparent for relatively high current injections of more than 500 pA and was not caused by depolarized plateauing with smaller AP amplitudes (Fig. 2A) as known from a truncation variant of *Scn1a* (Mattis et al. 2022), but was instead due to changes in the shape of the APs. At P20-23, APs within the train had a significantly slower rise time (0.31 ± 0.01 ms, n = 11 vs. 0.38 ± 0.02 ms, n = 14, p = 0.0013) - the only AP parameter already changed during the preseizure phase -, as well as a slower repolarization time (0.95 ± 0.06 ms, n = 11 vs. 1.19 ± 0.07 ms, n = 14, p = 0.0158) and an overall broader AP half width compared to the wt (0.31 ± 0.01 ms, n = 11 vs 0.40 ± 0.02 ms, n = 14, p = 0.0005) (Figure 2 G, H, I). During the stabilization phase at P37-40, the reduction in maximum firing frequency (Figure 2D) as well as other features in the AP waveform which reduced firing were found to be compensated or even reversed (Figure 2 G, H, I, Supplementary statistic table). fsIN of *Scn1a*^+/A1783V^ mice at P37-40 showed a faster AP rise time, faster repolarisation time and a reduced half width compared to wt fsIN. Additionally, phase plots for all three disease phases revealed in general smaller velocities of ΔV_m_ in fsIN of *Scn1a*^+/A1783V^ mice during the rising phase of the AP, particularly prominent at P20-P23 (Figure 2F).

**Figure 2.**
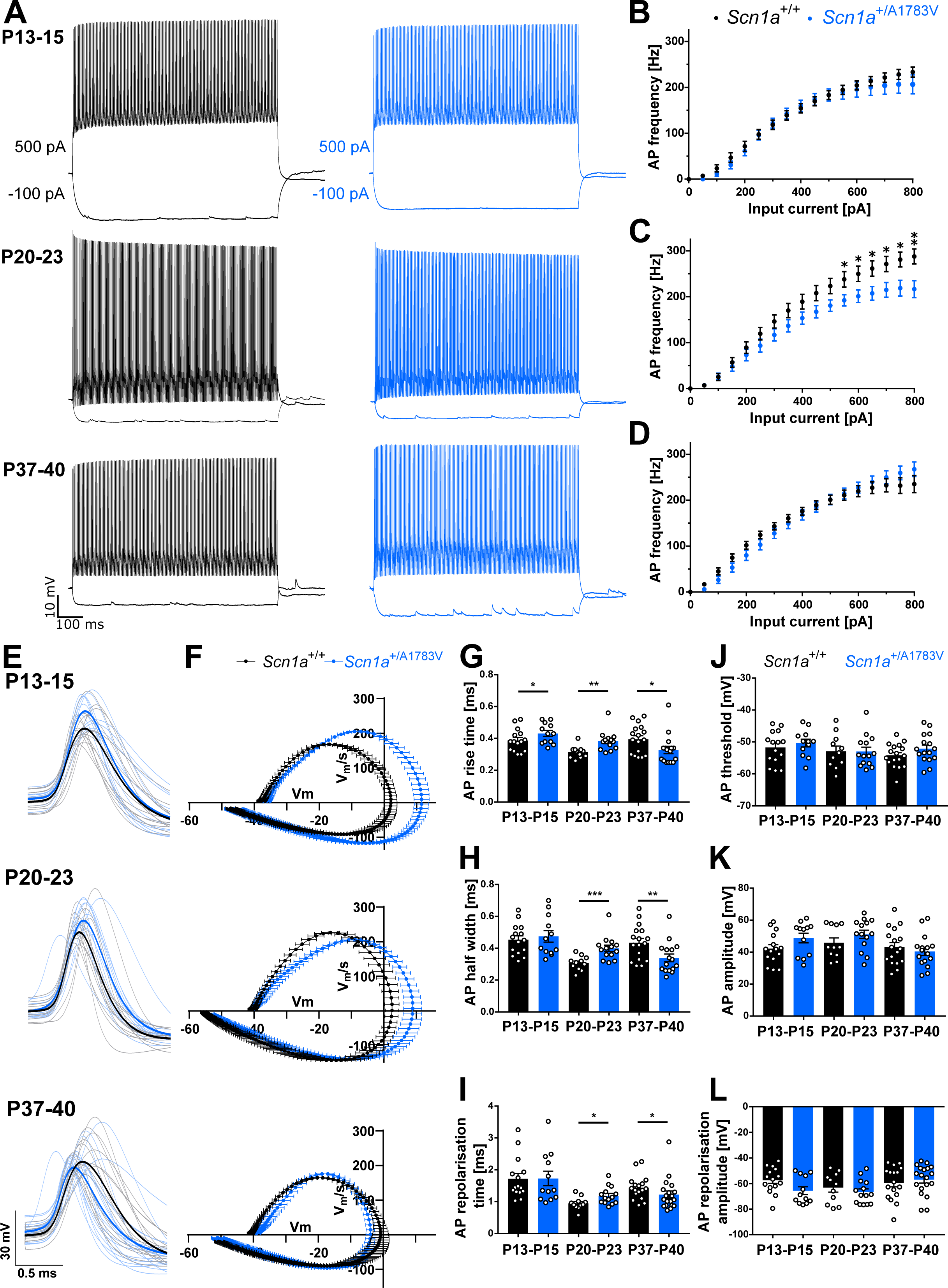
Alterations of CA1 fsIN AP properties and shape throughout the three disease phases. **A**: Representative AP trains at −100 pA and 500 pA current injection over a duration of 800 ms for fsIN of *Scn1a*^+/+^ (black) and *Scn1a*^+/A1783V^ (blue) mice before seizure onset (P13-15), at the high-seizure (P20-23) and the stabilization phase (P37-40). **B-D**: Frequency-current curves of fsIN throughout the three disease phases. AP frequencies were significantly reduced in *Scn1a*^+/A1783V^ mice for 550-800 pA current injection at P20-23 tested by multiple t-test. **E:** Waveforms of the last AP in train during an 800 ms long current injection of 150 pA in fsIN of *Scn1a*^+/+^ and *Scn1a*^+/A1783V^ brain slices before seizure onset, during the high frequency-seizure and the stabilization phase. Bold lines display the AP waveform mean per genotype, thin lines represent each datapoint. **F:** AP phase plots of the last AP in train during a 150 pA current injection of fsIN for *Scn1a*^+/+^ and *Scn1a*^+/A1783V^ brain slices before seizure onset, during high frequency-seizure and stabilization phase. **G-L**: Quantitative analysis of AP properties at the end of an AP train. APs from *Scn1a*^+/A1783V^ mice showed a significantly prolonged rise time, broader half width and prolonged repolarization time at P20-P23 while rise time, half width and repolarization time were significantly reduced at P37-P40 compared to *Scn1a*^+/+^ mice.

In contrast, we did not find such transient changes in excitability for regular-spiking interneurons (rsIN) in CA1 *stratum oriens*: There were neither differences in maximum firing frequency, single AP parameters nor in passive cellular properties between both genotypes at any of the three disease phases (Supplementary Figure 2, Supplementary statistic table). Similarly, somatic excitability of CA1 PCs was mostly unaffected; analysis revealed only mild alterations of the AP waveforms, as the first AP in a train displayed a broader half width (0.94 ± 0.06 ms, n = 8 vs. 1.13 ± 0.04 ms, n = 11, p = 0.0089) in the P20-23 age group (Supplementary Figure 3A-C, Supplementary statistic table). Altogether, these data demonstrate only mildly altered somatic excitability of interneurons which seems in stark contrast to the severe seizure phenotype of mice of the corresponding age group.

However, it remains unclear whether and how such mild alterations or other still unknown mechanisms impact network activity. To address this question, we next performed *in vitro* 2-photon calcium imaging in CA3 at P20-23. AAV-PHP.eB hSyn-gCamp7f was injected in the lateral ventricle of newborn mice. Consistent with the literature (Liu et al. 2020), *ex vivo* spontaneous network activity was most robust and reproducible in CA3 in our *in vitro* preparation. Recordings revealed a reduction of overall calcium-peak frequencies (0.26 ± 0.004 Hz, n = 271 vs. 0.24 Hz ± 0.002 Hz, n = 983, p = 0.00213) while the neuronal bursting activity was increased with significantly longer calcium-peak durations (3.08 ± 0.055 s, n = 271 vs. 4.12 ± 0.036 s, n = 983, p < 0.0001), larger peak area under the curve (AUC) (1.742 ± 0.465, n = 255 vs 4.10 ± 0.357, n = 926, p < 0.0001) and peak amplitudes (0.1917 ± 0.009 ΔF/F_0_, n = 271 vs. 0.23 ± 0.0.008 ΔF/F_0_, n = 983, p < 0.0001) as well as a higher maximal cross-correlation between pairs of neurons within brain slices of *Scn1a*^+/A1783V^ mice (0.05 ± 0.001, n = 5828 vs. 0.06 ± 0.0004, n = 56989, p < 0.0001), altogether indicating network hyperexcitability in CA3 (Supplementary Figure 4).

There is a clear discrepancy between the emerging network scale hyperexcitability and the mildly reduced somatic AP firing phenotype of fsIN during the phase of the highest seizure frequency. Furthermore, fsIN somatic excitability was completely normal at P13-15 (just before seizure onset). This contrasts with the transcriptome-based GO term analysis that suggests pathophysiological alterations occur at least as early as P16. Altogether, these data indicate that yet to be uncovered mechanisms distinct from reduced somatic excitability of fsIN may have to be considered as main drivers of initial pathogenesis.

### Loss of hippocampal inhibitory synaptic inputs precede seizures in *Scn1a*^+/A1783V^ mice and are uncoupled from somatic excitability of fsIN

As our snRNAseq analysis suggested early pathways alterations associated with synaptic structure and function, we next characterized inhibitory synaptic inputs in CA1 PCs of *Scn1a*^+/A1783V^ mice in comparison to wt. At P20-P23, spontaneous inhibitory postsynaptic current (sIPSC) amplitudes (−108.8 ± 7.19 pA, n = 25 vs. −80.1 ± 4.37 pA, n = 36, p = 0.0016) and the instantaneous frequency of sIPSCs were reduced (26.4 ± 3.54 Hz, n = 25 vs. 12.7 ± 1.38 Hz, n = 36, p = 0.0001). Similarly, miniature inhibitory postsynaptic current (mIPSC) frequencies were also reduced (16.27 ± 13.1 Hz, n = 19 vs. 6.798 ± 4.064 Hz, n = 19, p = 0.0018), although mIPSC amplitudes were slightly increased (−56.91 ± 2.96 pA, n = 19 vs. −65.43 ± 2.15 pA, n = 19, p = 0.0254) (Figure 3E, F, G, P, Q). By and large, reduced IPSC frequencies and amplitudes are consistent with previous studies in other Dravet mouse models (Stein et al. 2019; Uchino et al. 2021). At P13-15, mIPSC frequencies and amplitudes were unchanged between wt and *Scn1a*^+/A1783V^ cells (Figure 3N, O). However, at this early developmental time point, the sIPSC frequencies were significantly decreased (29.2 ± 3.54 Hz, n = 20 vs. 11.1 ± 1.81 Hz, n = 25, p = 0.0001) (Figure 3A, B, C). Additionally, sIPSCs were characterized by a slower decay as reflected by significantly increased time constants Δ_Decay_ for *Scn1a*^+/A1783V^ mice in comparison to wt animals (4.43 ± 0.18 ms, n = 21 vs. 5.71 ± 0.28 ms, n = 25, p = 0.0007) (Figure 3D). Although this slowing appeared to be transient and only present during the preseizure disease phase (Figure 3H), it brought up the question whether the reduced GABAergic transmission could in part derive from transcriptionally dysregulated GABAergic synaptogenesis resulting either in reduced numbers of inhibitory (most likely PVIN) synapses onto PCs or disrupted maturation of these synapses (see Discussion).

**Figure 3.**
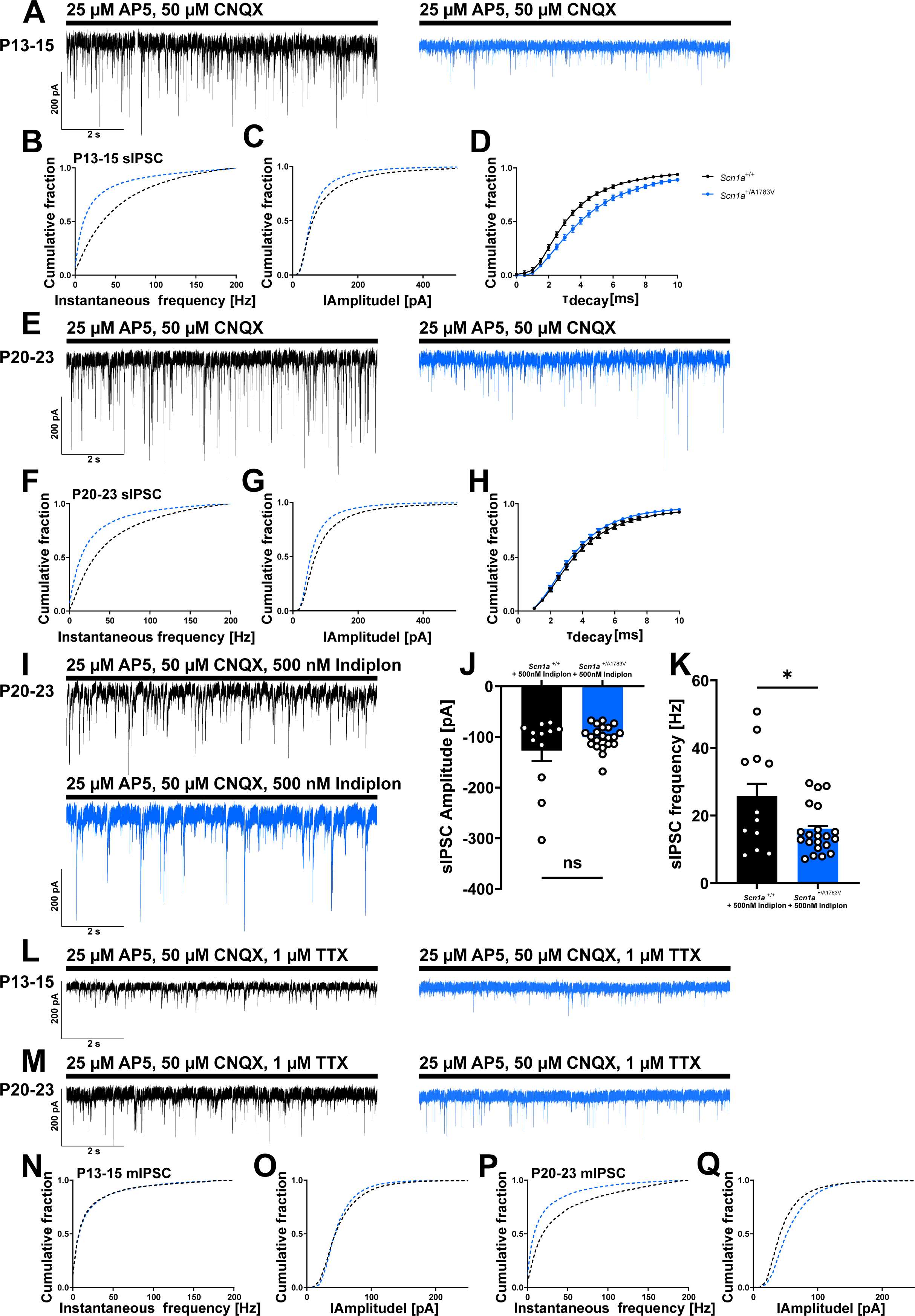
Inhibitory postsynaptic currents are impaired in CA1 pyramidal cells in *Scn1a*^+/A1783V^ mice before phenotypic seizure onset. **A, E**: Representative traces of spontaneous inhibitory postsynaptic currents (sIPSC)s recorded from CA1 PCs at P13-15 before seizure onset and P20-23 under application of AP5 and CNQX in *Scn1a*^+/+^ and *Scn1a*^+/A1783V^ mice. **B-D**: Cumulative fraction plots of event instantaneous frequency, amplitude, and time constant of event decay Δ_Decay_ at P13-15. sIPSC frequency and amplitude were significantly lower in *Scn1a*^+/A1783V^ mice and Δ_Decay_ was increased compared to *Scn1a*^+/+^ neurons. **F-H**: Cumulative fraction plots of event instantaneous frequency, amplitude, and time constant of event decay Δ_Decay_ at P20-23. sIPSC frequency and amplitude were significantly lower in *Scn1a*^+/A1783V^ mice. **I**: Representative traces of sIPSC recordings at P20-23 in CA1 PCs of *Scn1a*^+/+^ and *Scn1a*^+/A1783V^ mice after bath application of 500 nM indiplon. **J, K**: Quantitative analysis of sIPSC amplitude and frequency after indiplon bath application. 500 nM indiplon rescued sIPSC amplitude but not frequency in *Scn1a*^+/A1783V^ mice. **L, M:** Miniature inhibitory postsynaptic currents (mIPSCs) recorded from CA1 PCs in *Scn1a*^+/+^ and *Scn1a*^+/A1783V^ mice before and after seizure onset. **N-Q**: Cumulative plots displaying instantaneous event frequency and amplitude of mIPSCs.

To explore these questions at the transcriptional level, we first analyzed the time course of expression of gephyrin as a structural component of the postsynaptic density of inhibitory synapses in CA1 PCs (Craig et al. 1996). We separately clustered CA1 PCs without dataset integration for both time points which yielded separated clusters for *Scn1a*^+/A1783V^ cells at P20 identified as ventral and dorsal cell populations based on expression of ventral CA1 marker Dcn and dorsal CA1 marker Wfs1 (Cembrowski et al. 2016). While there was no change in expression of gephyrin at P16 (Supplementary Figure 5H), pseudotime analysis (see method section) surprisingly revealed an upregulation of gephyrin in CA1 PCs at P20 in the dorsal population, indicative of increased GABAergic synaptogenesis (Supplementary Figure 5I). To validate this, we immunohistochemically quantified the number of GABAergic synapses onto PCs in CA1. PV basket cell-derived perisomatic synapses were detected based on the presynaptic (PVIN synapse specific) marker Syt2. Consistent with the transcriptomic prediction of upregulated gephyrin, we indeed found an increased amount of perisomatic presynaptic terminals of PVINs in *Scn1a*^+/A783V^ mice in comparison to wt controls at P20 (21.56 ± 0.67 puncta/soma, n = 80 vs. 24.46 ± 1.14 puncta/soma, n = 79, p = 0.0292) (Supplementary Figure 5D - F). Similarly, an additional analysis of gephyrin puncta co-localizing with the axon initial segment marker ankyrin G revealed a substantially higher number of axo-axonic inhibitory synapses onto PCs (1.01 ± 0.07 puncta/µm, n = 43 vs. 1.42 ± 0.04 puncta/µm, n = 70, p = 0.0001) (Supplementary Figure 5A - C). These findings are not compatible with a loss of inhibitory synapses but rather represent compensatory mechanisms triggered by an increasingly emerging E/I imbalance.

To functionally test if a disrupted developmental switch from an immature to a mature GABA_A_ receptor subunit composition of PVIN synapses could explain loss of inhibitory input, we next recorded sIPSCs in CA1 PCs of brain slices derived from *Scn1a*^+/A1783V^ and wt animals in presence of indiplon (Figure 3I). Indiplon is a specific allosteric modulator of the α_1_-subunit containing GABA_A_ receptor, enriched in mature synapses of PV-positive fsIN projecting on PCs and as such enhances the amplitudes of PV basket cell (fsIN)-derived sIPSCs in presence of GABA (Petroski et al. 2006). Recordings from CA1 PCs in presence of indiplon mitigated the difference of sIPSC amplitudes between the wt and the *Scn1a*^+/A1783V^ condition (Figure 3J) whereas expectedly the sIPSC frequencies remained smaller in *Scn1a*^+/A1783V^ neurons (25.76 Hz ± 4.33, n = 12 vs. 16.06 Hz ± 1.54, p = 0.0165) (Figure 3K). These data indicate the presence of α_1_-subunit containing GABA_A_ receptors in *Scn1a*^+/A1783V^ PCs and are consistent with the establishment of an appropriate GABA receptor subunit composition at the postsynaptic side of inhibitory synapses in CA1 (see Discussion). This and the overall increased number of inhibitory synapses stands in contrast with our observations of less GABAergic inputs and therefore suggest a functional deficit of GABAergic interneurons and synapses caused by other still unknown pathomechanisms.

Altogether these data show that pathophysiological features of impaired GABAergic transmission precede seizure onset, develop before and alongside impaired somatic fsIN excitability and is accompanied by an emerging compensatory mechanisms. However, these changes in the early disease phases are not sufficient to counteract reduced inhibitory synaptic signaling.

### Identification of transcriptional changes in neuronal subpopulations contributing to epileptogenesis and network remodeling in the hippocampus of *Scn1a*^+/A1783V^ mice

Hyperexcitable networks that promote epileptic seizures can be derived from an altered interplay between excitatory and inhibitory neurons and are thought to be due to perturbed single cell and synaptic function (Jefferys 2010). To get insight into these processes at the transcriptomic level and further understand pathomechanisms involved in inhibitory dysfunction, we next investigated differential gene expression for inhibitory and excitatory neuron subpopulations at P16 and P20 using the Wilcoxon rank sum test. Mitochondrial genes were considered contaminations and were not taken into account. First level analysis revealed a relatively small number of genes changed at P16 (Figure 4A), confirming that disease pathogenesis is still in early stages at this time point. Only four days later, at P20, the number of differentially regulated genes had considerably increased in both excitatory and inhibitory cell populations (Figure 4B), in accordance with disease progression to phenotypic seizure onset. We next filtered for transcriptional determinants of neuronal excitability and synaptic connectivity by matching all genes (identified by Wilcoxon rank sum test disregarding p-values) with HUGO Gene Nomenclature Committee (HGNC) gene lists. These included curated lists for voltage-gated ion-channels, ligand-gated ion-channels, receptor-tyrosine kinases (RTKs), neurotrophins and neuregulins as well as cell adhesion molecules (CAMs) as all these molecule classes can majorly impact neuronal excitability. Corresponding heat maps were generated for CA1 and CA3 PCs as well as PV and SST interneuron clusters visualizing up- or down regulation. Also here, transcriptomic changes grew dynamically from initially few at P16 to then larger numbers of differentially regulated genes at P20 confirming that our dataset indeed mirrors the onset of epileptogenesis. These are likely to be capturing early disease mechanisms initiated before seizure onset, and secondary transcriptional alterations directly following first seizures. Within the class of CAMs predominantly genes of the neurexin, contactin and contactin-associated protein families were transcriptionally dysregulated (Figure 4E). For ligand-gated channels, there was an accumulated downregulation of NMDA-, AMPA- and kainate receptor subunit encoding genes in interneurons and CA1 PCs after seizure onset as well as prevalent dysregulation of GABA-receptor genes in all major neuron types reflecting seizure-dependent changes of synaptic plasticity in the postsynaptic cells (Figure 4D). Furthermore, we also found transcriptional downregulation of *Erbb4* and *Ntrk2* encoding Erb-B2 receptor tyrosine kinase 4 and tropomyosin receptor kinase B (TrkB receptor), respectively, in PVINs and CA1 PCs, as well as a general upregulation of the neurotrophin TrkB receptor ligand *Bdnf* across all four neuron populations. In the group of voltage-gated ion channels, predominantly potassium and calcium channel encoding genes were found dysregulated (Figure 4C). Amongst these genes, some were already downregulated at P16 including the potassium channel subunits *Kcnc1* in PVINs and *Kcnb2* in CA3 PCs, pointing towards early (“primary”) transcriptomic changes during epileptogenesis independent of (and not secondary to) seizures. After seizure onset, additional ion channel genes involved in membrane potential regulation (exemplified by *Kcnj3*, *Kcnh7* and *Kcnq3*) and repetitive neuronal firing (e.g. *Cacna1g* and *Cacna1h*) became dysregulated in excitatory neuron populations.

**Figure 4.**
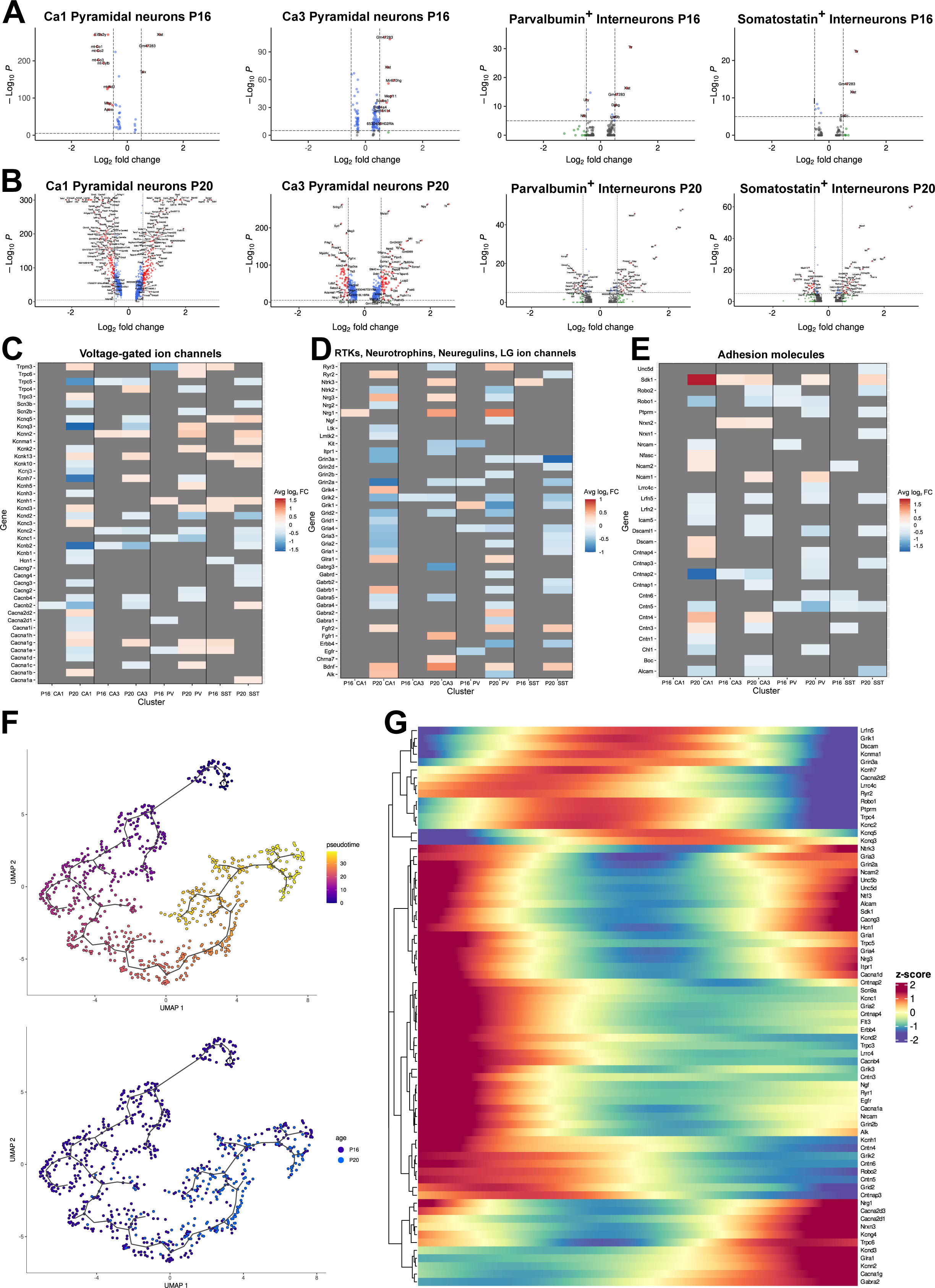
Pathologic alterations of gene expression of neuronal populations in hippocampus along the early phase of epileptogenesis. **A, B**: Volcano plots displaying significantly dysregulated genes in *Scn1a*^+/A1783V^ mice at P16 and P20 in the excitatory and inhibitory hippocampal neuron subtypes. Significantly dysregulated genes were identified by Wilcoxon rank sum test. Dashed lines represent cutoffs for average log_2_ fold change (FC) (│2^0.5^│) and adjusted p-value (p < 10^-5^). **C, D, E**: Heatmaps of dysregulated genes of selected gene families in excitatory and inhibitory neuron subtypes in the hippocampus at P16 and P20 in *Scn1a*^+/A1783V^ mice. Genes displayed in color show a >0.25 log_2_ FC change; the color code indicates the average log_2_ FC of these genes, gray boxes indicate log_2_ FC change <0.25. **F:** Umap representations of hippocampal PVINs of *Scn1a*^+/A1783V^ mice at P16 (purple) and P20 (blue) subclustered based on all dysregulated genes at P20 labeled as function of pseudotime and age using Monocle 3. **G**: Heatmap of genes describing the PVIN pseudotime trajectory with a cutoff of Moran’s i > 0.1 and a q-value > 0.05 (x-axis indicates pseudotime). Genes were clustered based on the pattern of change in expression along the pseudotime trajectory.

### Interneurons display a gradual transcriptomic trajectory during epileptogenesis

With differential expression testing, transcriptomic differences can only be assessed between clusters of cells but do not represent heterogenicity between gene expression profiles of individual cells. Therefore, this approach enabled us only to identify overarching differences between a physiological and a diseased network at two states to disease but failed to detect the first transcriptional changes that happen initially in few cells first affected in hippocampal epileptogenesis. It is conceivable that cells of a defined subtype will not progress simultaneously from a physiological towards a pathological state but rather follow a trajectory at a cell-specific pace. Based on this assumption, pseudotime trajectory analysis can be used in our data set to describe the cell-specific progress towards pathogenicity as well as to identify different cell states.

As the Na_V_1.1 pathogenic variant is thought to predominantly affect PVINs in the initial phase of epileptogenesis, we re-clustered all PVINs from *Scn1a*^+/A1783V^ mice of both age groups based on principal component analysis (PCA) restricted to the subset of genes found to be differentially expressed at P20 of our initial dataset analysis. The resulting UMAP representation from PCAs, which described at least four standard deviations, revealed a spatial trajectory of pseudotime (see method section, Figure 4F). Labeling for age revealed that some P16 interneurons co-clustered with the majority of PVINs at P20 while there were also some PVINs at P20 which co-clustered with P16 PVINs whereas other cells clustered in between. This indicates that transcriptomic changes develop gradually over time in individual PVINs. To identify genes progressively altered in their expression along pseudotime from a healthy to a pathological state, we used spatial autocorrelation via Moran’s statistics (Figure 4G). *Kcnc1*, *Kcnc2* and *Erbb4* were standing out amongst genes downregulated with increasing pseudotime and known to play key roles in specialized firing properties (Goldberg et al. 2005) and synaptic plasticity of PVINs (Chen et al. 2010). A similar analysis approach was taken for SST interneurons (Supplementary figure 2D, E) and CA1 PCs (data not shown). Whereas for SST interneurons the pseudotime trajectory calculation from P16 to P20 was successful and revealed, among others, progressive upregulation of potassium channel subunit encoding genes *Kcnq3* and *Kcnq5*, PCs did not cluster in a progressive trajectory but rather segregated into completely distinct clusters, indicating that most transcriptional changes in excitatory cells do not arise secondary to reduced GABAergic input in the preseizure phase but are rather triggered by the onset of first phenotypic seizures.

### Perisomatic CA1 synapses display ultrastructural alterations during early epileptogenesis

Another overrepresented group of differentially expressed genes were CAMs, mostly neurexins, contactins and contactin-associated proteins. Among these genes, we found presynaptic complex forming *Cntn5* and *Cntnap4* in PVINs, as well as postsynaptic *Nrcam* and *Chl1* in CA1 PCs to be downregulated in their expression at P20. At P16 in contrast, there was only a mild reduction of *Cntn5* in PVINs (Figure 5A, B). As these molecules are known to be binding partners at presynaptic terminals of perisomatic PVINs synapses and can affect synaptic morphology and consequently electrophysiologic synaptic transmission (Karayannis et al. 2014), we performed transmission electron microscopy (TEM) and imaged perisomatic symmetric synapses in CA1 *stratum pyramidale* (Figure 5C) at P15 and P20, and quantified synaptic length, width of the synaptic cleft as well as the number of presynaptic vesicles (Figure 5D-F). At P15 we found a significant reduction in the mean length of the postsynaptic density (PSD) (283.8 ± 10.28 nm, n = 78 vs. 237.9 ± 10.50 nm, n = 79, p = 0.0056) while the width of the synaptic cleft and the number of synaptic vesicles were unaltered between both genotypes. At P20 however, synaptic PSD length of wt mice had increased compared to P15 (283.8 ± 10.28 nm vs. 337.7 ± 12.49 nm), while there was no PSD length increase in *Scn1a*^+/A1783V^ mice (237.9 ± 10.50 nm vs. 241.6 ± 9.40 nm) resulting in even shorter perisomatic synapses in *Scn1a*^+/A1783V^ mice compared to the wt by P20 (337.7 ± 12.49 nm, n = 99 vs. 241.6 ± 9.40 nm n = 95, p <0.0001). Additionally, the width of the synaptic cleft was significantly wider in *Scn1a*^+/A1783V^ compared to the wt at P20 (11.3 ± 0.24 nm, n = 99 vs. 12.6 ± 0.22 nm, n = 95, p = 0.0011). Altogether these data suggest not only signs of a delayed synaptic maturation in *Scn1a*^+/A1783V^ mice but also an early pathologic structural development deficit that limits the efficiency of neurotransmitter transmission at GABAergic perisomatic synapses in CA1, consistent with reduced sIPSC amplitudes ((Karayannis et al. 2014); see Figure 3G and Discussion).

**Figure 5.**
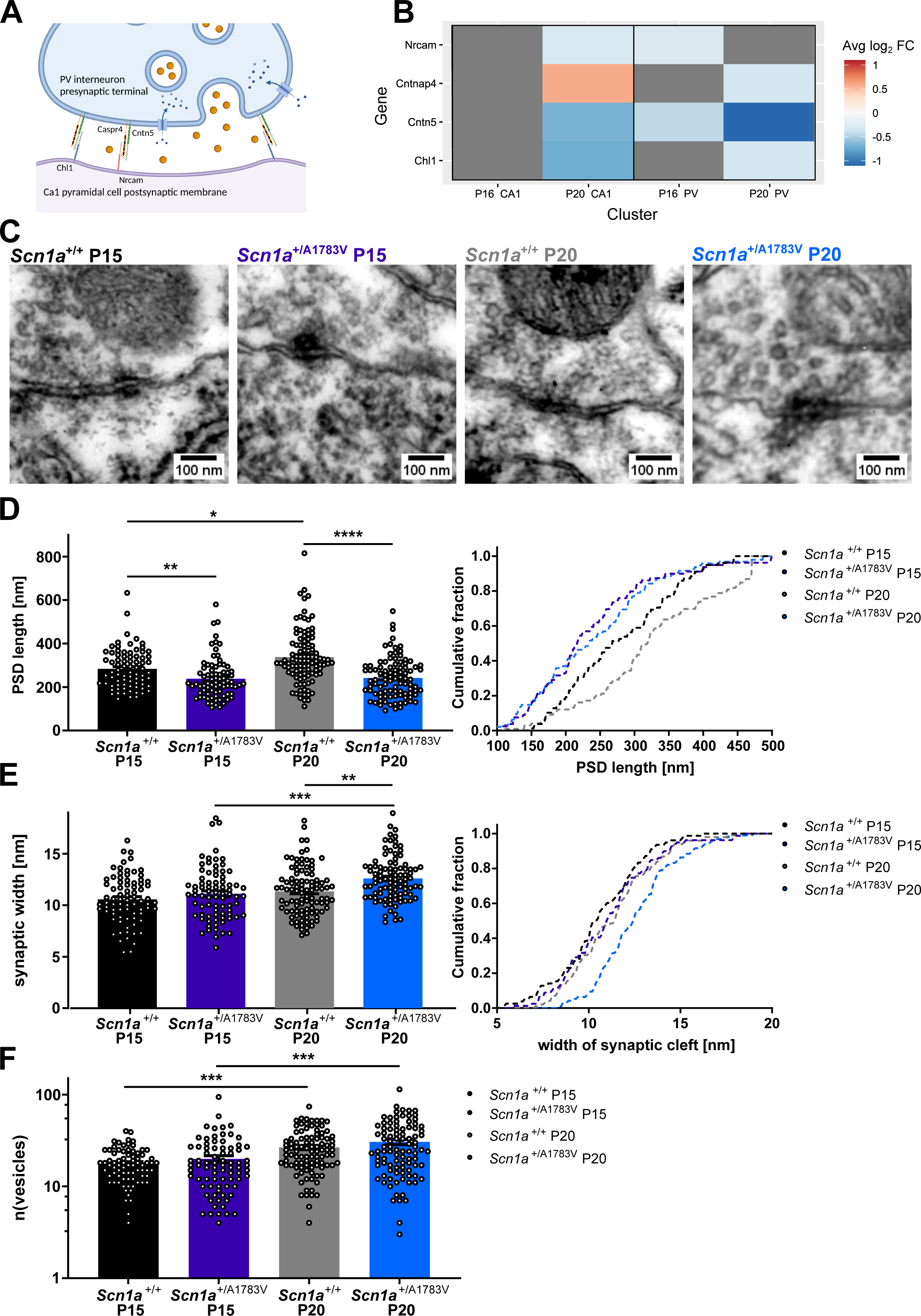
Transmission electron microscopy (TEM) of perisomatic synapses on CA1 neurons. **A**: Gene expression heatmap displaying dysregulated selected synaptic transmembrane CAMs in hippocampal neurons at P16 and P20. The color-bar indicated the magnitude of gene expression dysregulation in *Scn1a*^+/A1783V^ neurons compared to the wt. **B**: Representative images of perisomatic symmetric synapses of wt (*Scn1a*^+/+^) and *Scn1a*^+/A1783V^ mice at P15 and P20 in CA1 *str. pyramidale*. **C, D**: Cumulative fraction plots and scatter-dot plots of postsynaptic density length and synaptic width in EM pictures. Synaptic length was shortened in *Scn1a*^+/A1783V^ mice at P15 and P20. *Scn1a*^+/A1783V^ synapses at P20 also displayed a significantly enlarged width compared to wt (*Scn1a*^+/+^) at P20 and *Scn1a*^+/A1783V^ mice at P15. **E**: Number of presynaptic vesicles per imaged synapse. Vesicle numbers of both wt (*Scn1a*^+/+^) and *Scn1a*^+/A1783V^ mice increased significantly from P15 to P20.

Consequently, these data indicate that initially impaired interneuron dysfunction interferes with proper synapse development and maturation on the ultrastructural level, both of which have so far not been described in models of Dravet syndrome.

### Early hippocampal PV interneuron dysfunction is characterized by functional vulnerability of distal axonal and presynaptic compartments

While TEM had unraveled ultrastructural changes at perisomatic synapses, known from prior work on autism spectrum disorder (ASD) to promote progressive reduction of sIPSC amplitudes, the mechanisms underlying the frequency reduction of IPSCs, particularly before seizure onset, remained elusive. Therefore, taking advantage of the PVIN-specific E2 enhancer (Vormstein-Schneider et al. 2020) (Figure 6B), we injected AAV-PHP.eB E2-Chr2-mCherry in the lateral ventricle of newborn mice to test if reduction of GABAergic signaling was linked to impaired axonal function in hippocampal PVINs. Using the Polygon DMD pattern illuminator system, we performed optogenetic stimulation of transduced neurons in CA1 in acute brain slices with cell-compartment-specific stimulation in the field of view (instead of using full-field illumination) and recorded optogenetically stimulated inhibitory postsynaptic currents (oIPSC) in CA1 PCs. We applied series of five light pulses at 10 or 40 Hz and illuminated either PVIN somata leading to AP initiation at the axon initial segment, or the perisomatic region around the patched pyramidal cell’s soma causing AP initiation just at the presynaptic terminal (Figure 6A). While stimulating PVINs in wt mice revealed reliable oIPSCs at both timepoints (P13-P15 and P20-P23) for both illumination sites and only rarely produced oIPSC failures (Figure 6C-F), we found a significant reduction of successful oIPSCs when stimulating PVIN somata (10 Hz: 0.93 ± 0.02, n = 17 vs. 0.75 ± 0.06, n = 12, p = 0.0036; 40 Hz: 0.86 ± 0.04, n = 17 vs. 0.58 ± 0.07, n = 10, p = 0.0026) or presynaptic terminals (10 Hz: 0.91 ± 0.04, n = 14 vs. 0.70 ± 0.08, n = 8, p = 0.0135; 40 Hz. 0.85 ± 0.05, n = 13 vs. 0.53 ± 0.09, n = 10, p = 0.0068) of neurons of *Scn1a*^+/A1783V^ mice at P20-P23 (Figure 6I, J, K). At P13-P15, however, there was no significant reduction of oIPSC success rates yet when stimulating at the PVIN soma but rather a tendency for axonal dysfunction at 40 Hz stimulation (Figure 6G). Interestingly, at stimulation with 40 Hz restricted to perisomatic terminals yielded significantly lower success rates for oIPSCs recorded in PCs of *Scn1a*^+/A1783V^ mice (0.82 ± 0.04, n = 14 vs. 0.67 ± 0.06, n = 14, p = 0.0371) (Figure 6H, K) indicating enhanced vulnerability of the distal PVIN axon prior to seizure onset. Although impaired axonal function generally can be attributed to Na_V_1.1 LOF in PVINs of *Scn1a*^+/A1783V^ mice, subsequent transcriptomic alterations as part of epileptogenic processes likely amplify or contribute to this phenomenon. As demonstrated above by pseudotime analysis (Fig. 4 C, G) several genes (including *Kcnc1* and *Kcnc2; (Goldberg et al. 2005; Chen et al. 2021)*) involved in controlling excitability and synaptic plasticity of fsIN are downregulated with disease progression and could mechanistically explain the progression over time from distal to proximal compartments, particularly in light of stable *Scn1a* expression in PVINs at P16 and P20 (data not shown). It must be noted that for both timepoints and stimulation sites, we also found neurons in *Scn1a*^+/A1783V^ mice that transmitted oIPSCs reliably and comparably to those in wt mice, suggesting that not all PVINs are equally affected as shown above by pseudotime analysis of hippocampal PVINs.

**Figure 6.**
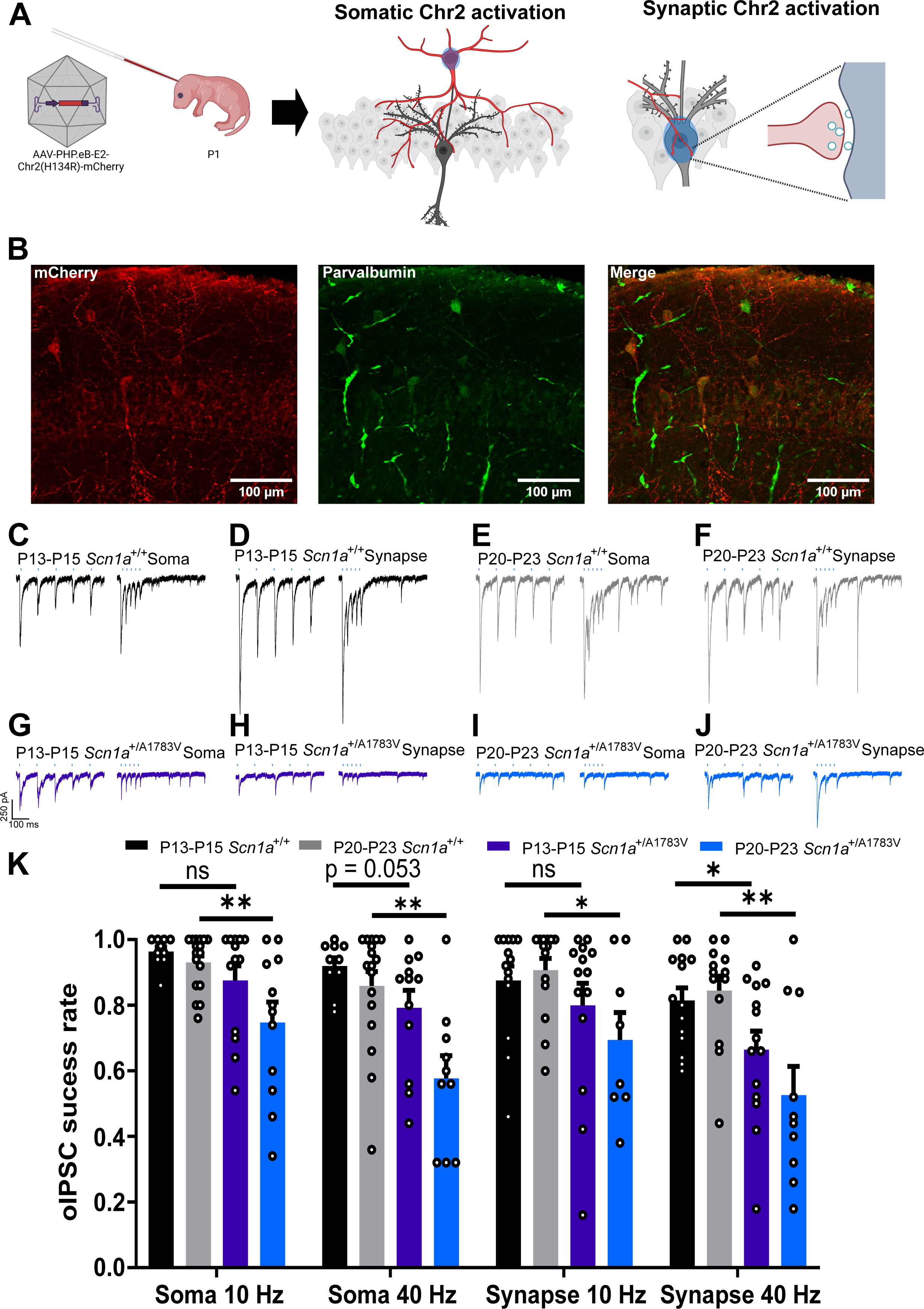
Cell-compartment specific optogenetic stimulation of PV interneurons in CA1. **A**: Schematic representation of AAV PHP.eB S5E2-ChR2-mCherry intracerebroventricular injection in postnatal mice and optogenetic stimulation of PVIN somata and presynaptic terminals. **B**: Representative IHC of post-hoc fixed brain slices with antibodies against mCherry (red) and PV (green). Hippocampal neurons expressing mcherry were all PV-positive while not all PVINs were transduced by the AAV. **C, E, G, I**: Representative oIPSCs recorded in postsynaptic PCs by 10 Hz (left) or 40 Hz (right) stimulation at the soma in wt (black, gray) and *Scn1a*^+/A1783V^ (purple, blue) neurons during the preseizure and high-frequency seizure phase. **D, F, H, J**: Representative oIPSCs recorded in postsynaptic PCs by 10 Hz (left) or 40 Hz (right) stimulation at the presynaptic terminal in wt and *Scn1a*^+/A1783V^ neurons before and after seizure onset. **K**: Quantification of successfully transmitted oIPSCs to the postsynaptic cell after PVIN somatic or presynaptic stimulation. *Scn1a*^+/A1783V^ PVINs displayed less successfully transmitted oIPSCs at P13-15 only when stimulating at the presynaptic terminal at 40 Hz, while *Scn1a*^+/A1783V^ PVINs at P20-23 showed significantly less successfully transmitted oIPSCs when stimulating at the soma or the synaptic terminal.

In short, restricted optogenetic stimulation revealed reduced oIPSC success rates in *Scn1a*^+/A1783V^ mice already before seizure onset which is more prominent at the distal axon/presynaptic terminal of hippocampal PVINs. This cellular phenotype intensified with seizure onset as we found both proximal (AIS/soma) as well as presynaptic compartments to be impaired at this later phase.

### Gene regulatory network modeling suggests a set of transcriptional regulators which drive differential gene expression throughout the course of epileptogenesis

While the analysis of our transcriptomic datasets revealed a multitude of primary and secondary transcriptomic changes during the process of epileptogenesis, the underlying regulatory cellular mechanisms are likely linked to altered activity of transcription factors (TF) (Valakh et al. 2023). To model potential TF (referred to as regulons from hereon) activity changes which could underlie differential gene expression identified in our transcriptomic data set during early epileptogenesis, we performed gene regulatory network (GRN) analysis for neuronal populations based on the hippocampal P20 snRNAseq data set. GRNs were built separately for inhibitory (Figure 7B) and excitatory (Figure 7D) cell clusters of each genotype. Mapping GRNs onto the previously annotated cell clusters resulted in remarkably specific representation of known neuronal regulons like *Lhx6* for interneurons derived from the medial ganglionic eminence (Alifragis et al. 2004) or *Foxp1* for CA1 PCs (Araujo et al. 2017), respectively (Figure 7A, C). As both GRNs reflected the expected marker regulon activity for both excitatory and inhibitory neuron types, we next compared neurons from wt and *Scn1a*^+/A1783V^ mice testing for differential regulon activity by Wilcoxon rank sum test (as had been done before for transcript counts) (Figure 7E-H). We then focused on all regulons predicted to affect expression of identified genes of interest dysregulated in *Scn1a*^+/A1783V^ mice, thereby aiming to connect the activity changes of regulons in the modeled GRN and the DEG of our snRNA data set. Matching genes and regulons this way, *Esrrg* was predicted to be less active in PVINs of *Scn1a*^+/A1783V^ mice, thereby controlling downregulation of potassium channel subunit encoding *Kcnc1*, sodium channel modifier *Fgf12*, *Ntrk2*, *Erbb4* and *Cntnap4* amongst others. Likewise, in excitatory neurons we found *Thra_extended* and *Klf3_extended* significantly less active in *Scn1a*^+/A1783V^ CA1 and CA3, thereby downregulating expression of *Kcnq3*, *Nrcam* and *Chl1*. In conclusion, GRN modeling highlighted a set of transcriptional regulators putatively underlying epileptogenesis in *Scn1a*^+/A1783V^ mice. These predictions may enable further dissection of epileptogenic mechanisms in follow up studies and may promote future identification of common regulatory networks among different forms of hippocampal epilepsy.

**Figure 7.**
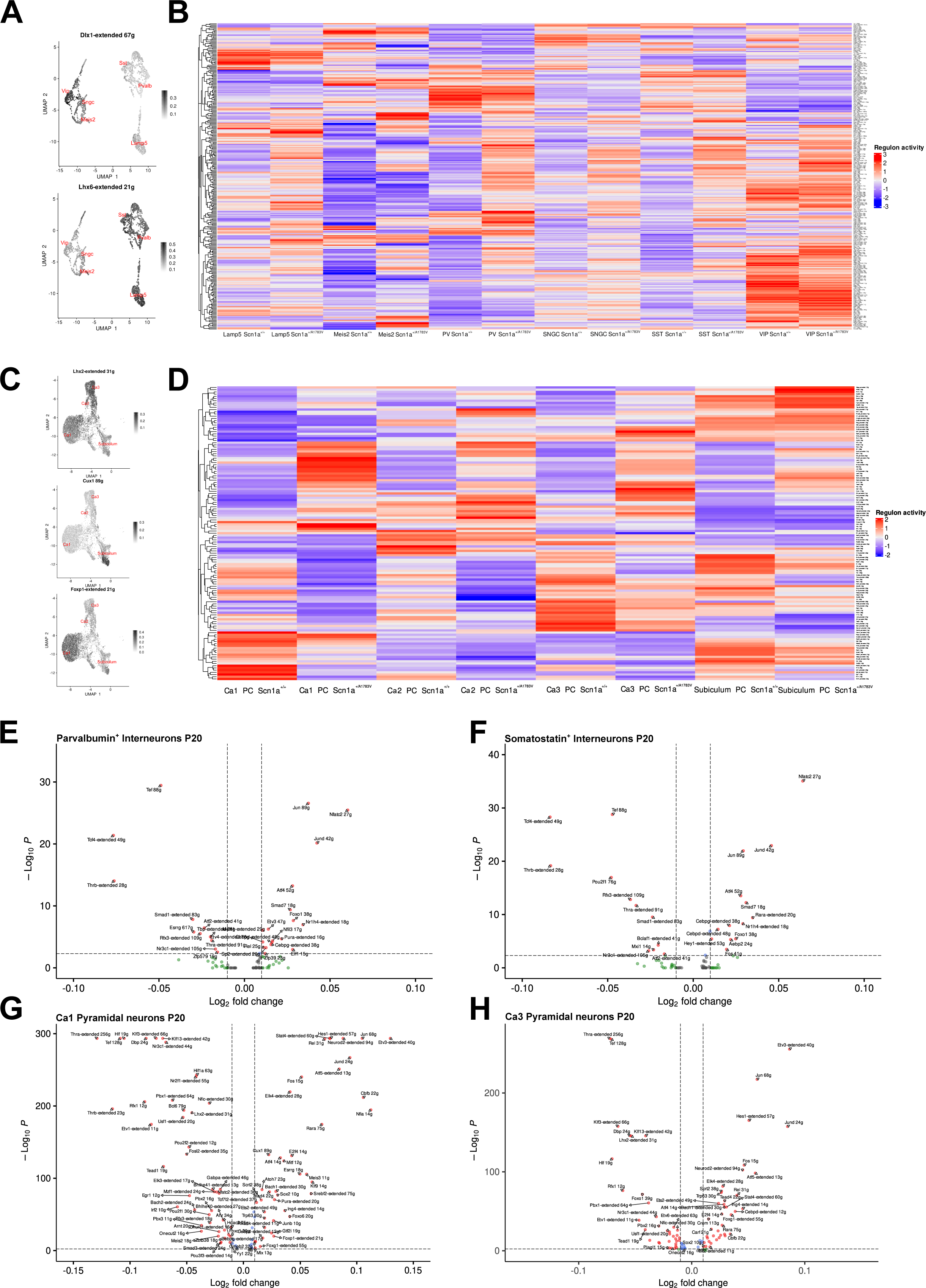
Modeling of a gene regulatory network to predict transcription factor activity in wt and *Scn1a*^+/A1783V^ mice at P20. **A**: Umap feature plot of TF linked GRN activity for marker TFs *Dlx1* and *Lhx6* in hippocampal interneurons indicated by color intensity. **B**: Heatmap of predicted activity of TF in a simulated GRN of hippocampal interneurons sorted by cell type and genotype. **C**: Umap feature plot of TF linked GRN activity for marker TFs *Lhx2*, *Cux1* and *Foxp1* in hippocampal PCs indicated by color intensity. **D**: Heatmap of predicted activity of TF in a simulated GRN of hippocampal PCs sorted by cell type and genotype. **E-H**: Volcano plots displaying significantly dysregulated TF activity in *Scn1a*^+/A1783V^ mice at P16 and P20 in excitatory and inhibitory hippocampal neuron subtypes. Significantly dysregulated TF activity was identified by Wilcoxon ranked test. Dashed lines represent cutoffs for average log_2_ FC (│2^0.01^│) and adjusted p-value (p < 10^-5^).

## Discussion

Prior work throughout the last two decades identified interneuron dysfunction as a hallmark of DS and Na_V_1.1 loss-of-function as the most common underlying molecular mechanism. The disease emerges during an early developmental time window of ongoing circuit maturation and enhanced synaptic plasticity. In mice this time course coincides with increasing Na_V_1.1 expression levels in cortex and hippocampus by the end of the second week of age (Cheah et al. 2013; Liang et al. 2021). Proper establishment of neuronal network connectivity and E/I balance depend on tightly regulated cell-intrinsic (e.g. Dehorter et al. 2015) and intercellular communication (e.g. Li et al. 2011) and voltage-gated ion channels have been demonstrated to be critically involved in such processes on multiple levels depending on their temporal, spatial and neuron subtype expression patterns (Gu et al. 2018; Schulz et al. 2008). We hypothesized that altered Na_V_1.1- and associated interneuron dysfunction will set off a cascade of transcriptional dysregulation representing the molecular basis of disease pathogenesis (including epileptogenic processes triggered by the *Scn1a* mutation) culminating in seizures and neurodevelopmental symptoms and being distinct from later stage secondary astrogliotic and inflammatory processes which have previously been reported (Miljanovic et al. 2021; Martín-Suárez et al. 2020; Hawkins et al. 2019; Alonso Gómez et al. 2018). To address this rationale, we performed single-nuclei RNA-sequencing in a mouse model of DS with single-cell resolution just before and shortly after seizure onset and in comparison to healthy littermate controls. First-pass GSEA analysis indicated transcriptional dysregulation of neurodevelopmental processes including synaptic signaling and channel activity in multiple neuron types clearly before seizure onset. Although somatic firing of CA1 fsIN in brain slices from *Scn1a*^+/A1783V^ mice was normal at this time point, GABAergic transmission to CA1 PCs was already markedly reduced. This discrepancy between somatic excitability and synaptic transmission by itself is a novel aspect of early IN pathology in DS which is not in conflict with prior studies as these have not consistently investigated IPSCs in parallel to somatic IN firing rates at the preseizure phase. Particularly, a recent study in *Scn1a*^+/-^ mice (Kaneko et al. 2022) demonstrated normalization of somatic spiking but persisting inhibitory synaptic transmission impairment in the stabilization phase, indicating that altered synaptic transmission does not exclusively depend on somatic properties of interneurons. However, the underlying mechanisms remained largely unknown. As pathological development progresses and seizures emerge, early transcriptomic alterations increasingly aggravate and a multitude of additional dysregulations occur in many cell types. At the same time, in contrast to truncation models of DS, in which fsINs cannot sustain prolonged high-frequency firing at the soma in cortex and hippocampus (Ogiwara et al. 2007; Favero et al. 2018; Goff und Goldberg 2019), we only observed a mild transient reduction of maximal firing rates and broadening of the somatic AP waveform in CA1 fsIN. This somatic hypoexcitability, contrary to previous findings (Almog et al. 2021), was not present in rsINs. We argue that this discrepancy is due to the different genetic background of the studied Dravet mouse line, since Dravet mouse models on a pure B6 background carry an additional intronic deletion mutation in *Gabra2* which aggravates the epileptic phenotype (Hawkins et al. 2021). Therefore, the mixed S129/Bl6 genetic background of *Scn1a*^+/A1783V^ mice used in our study reflects the genetic condition of DS patients more accurately and ameliorates the severity of IN dysfunction to levels comparable with the *Scn1a*^+/R1407X^ DS mouse model (Rubinstein et al. 2015).

As the primary mechanism of developmental hippocampal PVIN dysfunction, we identified a lack in IPSC generation in *Scn1a*^+/A1783V^ mice. Given the rather unremarkable somatic firing properties of PVINs and increased numbers of axo-axonic and perisomatic inhibitory synapses onto CA1 PCs in *Scn1a*^+/A1783V^ mice, we reasoned that the IPSC reduction most likely may originate from axonal dysfunction and/or altered synaptic transmission: GSEA and DEG analysis in PVINs indeed revealed downregulation of genes involved in axonal development and function as well as in assembly of GABAergic synapses. This hypothesis was experimentally explored by optogenetic stimulation of PVIN axonal and presynaptic compartments in CA1 revealing reduced success rates of GABAergic oIPSC. In contrast to prior work in cortical brain slices of the *Scn1a*^+/R1407X^ mouse model which had shown decreasing amplitudes of repetitively evoked inhibitory currents after somatic but not axonal whole field optogenetic stimulation, suggesting primary vulnerability at the proximal axon initial segment (Kaneko et al. 2022), we found that also the distal axonal compartment including the inhibitory presynapse is vulnerable to Na_V_1.1 LOF, well before somatic firing rates are affected. Nevertheless, the two studies were conducted in different Dravet mouse models with different primary mechanisms of disease: Truncation variants cause a reduced sodium conductance due to lower Na_V_1.1 channel density in the neuronal membrane while p.(Ala1783Val) alters voltage-dependence of activation and slow inactivation of Na_V_1.1 (Layer et al. 2021) reflected by increased AP rise times and smaller velocities of ΔV_m_ in our whole cell recordings of CA1 fsIN. Yet, experimental evidence of dysfunction distal to the somatic compartment in both mouse models shifts the focus of the primary pathomechanism of Dravet syndrome from reduced somatic firing rates towards a predominant axonopathy and synaptopathy. In support of this interpretation, we found dysregulation of several genes that likely promote interneuron dysfunction in *Scn1a*^+/A1783V^ mice: An early reduced expression of potassium channel encoding genes *Kcnc1* and *Kcnc2* in *Scn1a*^+/A1783V^ mice stood out from our transcriptomic dataset in PVINs and was already present during the initial phase of epileptogenesis before seizure onset. These channels are important in the regulation of axonal and presynaptic signaling in fsIN (Goldberg et al. 2005), are required for high-frequency AP firing and their expression directly correlates with the expression of sodium channels, particularly Na_V_1.1 (Gu et al. 2018). As proteomic bulk analysis in a genetic mouse model with reduced neddylation in PVINs leading to less Na_V_1.1 expression also displayed a reduction in K_V_3.1 protein levels (Chen et al. 2021), downregulation of K_V_3 potassium channels in interneurons might be a common secondary mechanism contributing to axonopathy in *Scn1a* LOF models.

Although the analysis of somatic firing rates remains an important parameter to evaluate neuronal dysfunction in models of DS, our data show that the loss of GABAergic synaptic transmission is an early key pathomechanism in *Scn1a*^+/A1783V^ mice developing independently from reduced somatic AP firing. This reduction of sIPSC amplitudes and frequencies, which we found to be present already in the second week of age, suggested either a reduced number of APs reaching the presynaptic GABAergic synapse, an overall lack of PV-PC synapses or alterations in postsynaptic GABA receptor density and composition and therefore altered integration of GABAergic inputs. As we found an overall increased number of inhibitory synapses at the high-seizure frequency phase (Supplementary figure 5), slower decay kinetics of sIPSCs in CA1 PCs particularly in the preseizure disease phase as well as shorter PSDs of symmetric (inhibitory) perisomatic synapses onto CA1 PCs, we propose that the development of PVIN synapses from an immature to a mature state is impaired in the *Scn1a*^+/A1783V^ mouse model. While mature PV-positive basket cell synapses are characterized by enrichment of GABA_A_ receptor α_1_-subunits, which are associated with fast chloride kinetics, immature PV-positive basket cells highly express α_2_-subunits and mediate IPSCs with smaller amplitudes and slower kinetics (Klausberger et al. 2002; Doischer et al. 2008; Nomura et al. 2019). Additionally, other interneuron subtypes including PV-negative basket cells but also PV-positive axo-axonic cells mediate inhibitory transmission via α_2_-subunit containing GABA_A_ receptors (Booker and Vida 2018). As our snRNA seq data did neither reveal dysregulated GABA receptor α_1_ nor α_2_-subunit expression in PCs at P16 nor P20, altered GABA receptor composition at the PVIN postsynaptic density (PSD) of PCs seemed unlikely to account for reduced synaptic transmission. This assumption was strengthened by our sIPSC measurements under application of the allosteric GABA_A1_ receptor modulator indiplon which revealed pharmaco-sensitivity of CA1 GABAergic synapses in *Scn1a*^+/A1783V^ mice at P20 pointing towards matured GABA receptor compositions localized to the PSD of perisomatic PVIN synapses. Interestingly, the increase of the sIPSC amplitudes under Indiplon application seemed more pronounced for neurons of *Scn1a*^+/A1783V^ mice than for wt controls which could be attributed to the increased numbers of inhibitory synapses at this age, most likely demonstrating an early compensatory mechanism to counteract network hyperexcitability and seizures. Nevertheless, we can’t completely rule out that the postsynaptic GABA receptor composition at PVIN synapses is altered in *Scn1a*^+/A1783V^ mice impacting postsynaptic integration of GABA signaling as we found a downregulation of *Gabra4* and *Gabra5* transcripts in CA1 PCs after seizure onset which might have an influence on the function of PVIN GABAergic synapses.

While these data so far did not support the hypothesis of a maturational deficit of PVIN synapses, ultrastructural changes of putative perisomatic PVIN synapses in CA1 strongly pointed in this direction and were consistent with smaller IPSC amplitudes. Our analysis of DEG in PVINs and PCs links progressively shorter and wider synaptic clefts identified by TEM in *Scn1a*^+/A1783V^ mice to underlying transcriptomic alterations, i.e. reduced expression of synaptic transmembrane CAMs subsequently leading to the reduction of sIPSC amplitudes. It was previously shown that the reduction of presynaptic contactins *Cntnap4* and *Cntn5*, which form an anchoring complex in the synaptic cleft with postsynaptic molecules *Nrcam* and *Chl1*, results in a loss of inhibitory boutons (Ashrafi et al. 2014) or ultrastructural alterations with reduced IPSC amplitudes at cortical perisomatic PVIN synapses similar to our findings in this study (Karayannis et al. 2014). We suggest that psychiatric comorbidities present in *Scn1a*^+/A1783V^ Dravet mouse model (Ricobaraza et al. 2019) could at least partly be linked to the synaptic ultrastructural alterations and reduction of postsynaptic IPSCs. Indeed, although most of these comorbidities are thought to arise from frontal cortical dysfunction (Xu et al. 2019; Ferguson und Gao 2018), the cellular origin of PVINs as well as their role in these circuits, providing local feedback and feedforward inhibition, is comparable between hippocampus and cortex (Nahar et al. 2021). Importantly, mutations in genes encoding for these CAMs, have been discussed in the clinical context as potential risk factors for the development of schizophrenia (Chen et al. 2005; Kim et al. 2009) and ASD (Li et al. 2016; Marui et al. 2009; Karayannis et al. 2014). As expression (Liakath-Ali and Südhof 2021) and cleavage (Suzuki et al. 2012; Peixoto et al. 2012) of different CAMs can be influenced by intrinsic neuronal activity and intracellular signaling pathways (Jiang et al. 2021), it seems conceivable that reduced axonal excitability of PVINs in the initial phase of epileptogenesis modulates the expression of CAMs and subsequently results in impaired synaptic ultrastructure.

Moreover, we found noteworthy transcriptomic alterations in hippocampal PVINs linking epileptogenesis to comorbidities in DS: Downregulated *Erbb4* was previously shown in knock-out mice to be crucial for PVIN maturation (Golub et al. 2004; Chen et al. 2010; Wang et al. 2018), and *Erbb4* deletion restricted to PVINs resulted in higher seizure susceptibility (Li et al. 2011). Regarding its ligands, *Nrg3* (Müller et al. 2018) expression was found to be reduced in PVINs while *Nrg1* levels were increased. Further, transcription of *Bdnf* signaling was dysregulated in excitatory and inhibitory neuron types in the CA region: Increased levels of *Bdnf* exacerbate epileptogenesis (He et al. 2004) while a chronic activation of its receptor Trkb, encoded by *Ntrk2*, which we also found to be downregulated in PVINs, can restore IPSC reduction and reduce seizure frequency in *Scn1a*^+/A1783V^ mice (Gu et al. 2022). All these alterations can be connected to deficits in motor development and memory consolidation which are present in *Scn1a*^+/A1783V^ mice (Ricobaraza et al. 2019; Reiber et al. 2022). Another hit with potential relevance for PVINs pathology was the reduction of *Fgf12* which has recently been reported to functionally alter the kinetics of sodium channels Na_V_1.2 (Wildburger et al. 2015) and Na_V_1.6 (Wang et al. 2011) and to cause DEE when disrupted by a missense variant or duplication variant (Seiffert et al. 2022). While a direct interaction between Na_V_1.1 and Fgf12 has not been reported, the modulation of intrinsic PVIN excitability by Fgf12 via Na_V_1.6 has to be considered in this context.

Even though the dysfunction of fsINs is most likely the primary driver of hippocampal epileptogenesis, snRNAseq at P20 revealed also secondary alterations in excitatory cell clusters that might contribute to network hyperexcitability: In contrast to *Scn1a*^+/R1407^ mice, in which the dentate gyrus was recently identified to be disinhibited and to promote hippocampal seizure generation (Mattis et al. 2022), we found a higher number of dysregulated genes and pathways in hippocampal CA1 and CA3 areas. Specifically, the downregulation of potassium channel encoding genes *Kcnj3*, *Kcnh7*, *Kcnq3* and *Kcnq5* in PCs suggested facilitated bursting behavior relevant for seizure generation in *Scn1a*^+/A1783V^ mice. The increased hippocampal network excitability was confirmed experimentally by two-photon calcium imaging in CA3, where we found prolonged neuronal bursting and increased network synchronicity within the *str. pyramidale* which spotlights the relevance of the *cornu ammonis* areas for the generation of hippocampal hyperactivity in *Scn1a*^+/A1783V^ mice. As *Scn1a* LOF variants can also have direct effects on early PC intrinsic excitability (Almog et al. 2021) and excitatory signaling (Jones et al. 2022), it might even be possible that the identified early transcriptomic changes in PCs lead to a parallel increased level of excitation that further promotes a pathologic shift of E/I balance. These early alterations in PCs, likely facilitating hippocampal seizure generation, might prove to become useful targets to develop novel therapeutic strategies targeting pathophysiologic remodeling in DS or other forms of hippocampal epilepsy. As we found not only potential pathologic alterations but also transcriptional changes which rather counteract network overexcitability at P20, - e.g. increase of GABAergic synapses and the reduction of glutamate receptor gene transcripts in excitatory cells which was also generally found in bulk proteomics in the same mouse line before (Miljanovic et al. 2021) - it becomes apparent that network compensation starts already with or before the first seizures and not once seizure occurrence declines and becomes more rare in *Scn1a*^+/A1783V^ mice around P40. Even further, all secondary changes found after seizure onset might even be considered as compensatory mechanisms toward reaching a new stable network state with occasional occurrence of seizures.

While the snRNA dataset presented in this study is restricted to a single mouse model of DS, it can be assumed that some pathway alterations are not specific to *Scn1a*^+/A1783V^ mice but are instead common between different hippocampal epilepsy models characterized by similar seizure types. Such shared pathomechanisms might include altered activity at the level of TFs. When exploring the modeled GRNs built from our hippocampal RNA seq dataset, some TFs predicted to be less active across excitatory and inhibitory neuron types point towards this concept: Recent studies have shown that in rodent models of acquired epilepsy by systemic kainic acid or pilocarpine injection a lower expression of TFs *Hlf*, *Tef* and *Dbp* can be found (Rambousek et al. 2020), while a knock-out of TF *Hlf* in a *Scn2a* mouse model increases seizure frequency and mortality rate (Hawkins and Kearney 2016). Furthermore, a triple homozygous knock-out mouse model of *Hlf*, *Tef* and *Dbp* - all these TFs were predicted to be less active in our TF model of hippocampal CA1 and CA3 PCs - led to network hyperexcitability and changes in homeostatic plasticity *in vitro* as well as to spontaneous seizures and premature death *in vivo* (Valakh et al. 2023). This supports the idea that there are common mechanisms underlying hippocampal epileptogenesis once the balance of excitation and inhibition has been disturbed by either Na_V_1.1 dysfunction like in Dravet syndrome or by other symptomatic or genetic causes.

Taken together, whereas the concept of epileptogenesis is well accepted for acquired temporal lobe epilepsy (Becker 2018), it has not been clearly established for genetic epilepsies, whether a disease-causing genetic variant is sufficient by itself to cause the full phenotypic extent of the pathology, or if additional epileptogenic processes and their interference with proper neurodevelopment are required. Our study provides a single-cell and time-resolved transcriptomic data set in a mouse model of DS spanning the critical time window of symptom onset and reveals strong evidence for an early hippocampal epileptogenetic process in DS – beyond the currently existing concepts of disease pathogenesis – which very likely not only significantly contributes to the epileptic phenotype but also to the emergence of neurodevelopmental symptoms. While the somatic excitability of hippocampal fsINs was only mildly and transiently affected by Na_V_1.1 LOF, we found evidence that the distal axonal and presynaptic compartments are particularly functionally vulnerable promoting reduced GABAergic transmission. Subsequently, initial interneuron dysfunction results in shifted E/I balance, leads to alterations in the numbers and the structure of GABAergic synapses and affects a multitude of further signaling pathways in excitatory and inhibitory cell populations linked to pathomechanisms of comorbidities. Our dataset will serve as a substantial resource for the epilepsy- and neurodevelopmental research community and will enable future studies of pathologic pathways and exploration of novel therapeutic concepts in all major hippocampal cell types of DS.

## Methods

### Animals

All conducted animal experiments were reviewed and approved by the local Animal Care and Use Committee (Regierungspraesidium Tübingen, Tübingen, Germany). *Scn1a*^+/A1783V^ mice were created by crossing heterozygous male B6(Cg)-*Scn1a^tm1.1Dsf^*/J and female 129S1/Sv-*Hprt1^tm1(CAG-cre)Mnn^*/J mice. Offspring were weaned after 23 days to reduce stress in *Scn1a*^+/A1783V^ mice during the high seizure frequency phase of the disease and additionally fed with milk powder to reduce weight loss due to seizures. Spontaneously occurring seizures were not medicated to not influence epileptogenesis. All animals bred for the purpose of this study were used for experiments before P40 to reduce the burden caused by the genetic variant. From P18 onwards animals were monitored regularly by the experimenters and experiments were discontinued if the level of phenotypic burden surpassed a predetermined score.

### Viral injections

Viral injections were performed postnatally at P1-2 with in-house produced according to or commercially available viral vectors AAV-php.eB-hSyn-gCamp7f (Addgene Plasmid #104488) or AAV-php.eB-S5E2-ChR2-mCherry (Addgene Plasmid #135634) as followed: After applying Emla cream (2.5% Lidocaine, 2.5% Pilocaine) to the skin covering the skull, newborn mouse pups were briefly immobilized by hypothermia for up to one minute until movement arrest. Subsequently, viral vectors were bilaterally injected intracerebroventricularly using a beveled glass pipette with a volume of 1.5 µl virus mixed with Fast Green dye (virus titer: 1×10^13 vg/ml) per hemisphere. Subsequently, mice were briefly warmed up under a red light and returned to the home cage.

### Single nuclei RNA sequencing sample preparation

Sample extraction for single nuclei RNA sequencing was performed at P16 and P20 from three wt and three *Scn1a*^+/A1783V^ mice by dissection of one brain hemisphere per mouse resecting the hippocampus. Hippocampal tissue was collected and frozen covered in homogenization buffer (Sucrose 250 mM, KCl, MgCl_2_ 5 mM, Tris pH 8.0 10 mM, DTT 0.001 mM, cOmplete™ 1x (Roche, Switzerland), Triton X-100 1µl/ml, RNAsin 0.4 U/µl) at −80°C until the day of library preparation. On the day of preparation, samples were thawed on ice, homogenized by 10-15 up- and downward strokes with a pistil free from RNAses. Homogenates were centrifuged for eight minutes at 1000x g and the supernatant was discarded. Pellets were resuspended in homogenization buffer and passed through a 40 µm strainer (Greiner Bio-One, Germany). Flow through was adjusted to a final Iodixanol concentration of 25% and layered on top of a gradient with 29% Iodixanol (Stemcell Technologies). After spinning the samples for 20 minutes at 13500x g the supernatant was carefully removed without disrupting the remaining purified nuclei pellet. In a final step, nuclei were resuspended in PBS with 1% BSA (Sigma-Aldrich) containing 0.4 U/ml RNAse inhibitor (New England Biolabs) and were immediately transported to the sequencing facility on ice for further processing.

### 10x Genomics single-cell RNA sequencing

Single nuclei suspension concentration was determined by automatic cell counting (DeNovix CellDrop, DE, USA) using an AO/PI viability assay and counting nuclei as dead cells. Gene expression libraries were generated using the 10x Chromium Next gel beads-in-emulsion (GEM) Single Cell 3’ Reagent Kit v3.1 (10x Genomics, CA, USA) according to manufacturer’s instructions. In brief, up 20,000 nuclei originating from mouse brain together with barcoded Single Cell 3’ v3.1 Gel Beads and Partitioning Oil were loaded on the Chromium Next GEM Chip G, which was subsequently run on the Chromium Controller (10x Genomics, CA, USA) to partition nuclei into GEMs. Afterwards, the Gel Beads were dissolved, primers were released, and reverse transcription of poly-adenylated mRNA using the poly(dT) primers occurred within the GEMs and resulted in cDNA with GEM-specific barcodes and transcript-specific unique molecular identifiers (UMIs). Silane magnetic beads were applied to purify the cDNA from the post GEM-RT reaction mixture. In the next step, full-length cDNA was amplified by PCR to produce sufficient mass for library construction. Enzymatic fragmentation, and size selection were done to optimize the cDNA amplicons. Full-length cDNA was end-repaired, extended with 3′ A-overhangs, and ligated to adapters. P5 and P7 sequences, as well as sample indices (Chromium i7 Multiplex kit, 10x Genomics, CA, USA), were added during the final PCR amplification step. The fragment size of the final libraries was determined using the Bioanalyzer High-Sensitivity DNA Kit (Agilent, CA, USA). Library concentration was determined using the Qubit dsDNA HS Assay Kit (Thermo Fisher Scientific, MA, USA). Single Cell 3’ Gene Expression libraries were pooled and paired-end-sequenced on the Illumina NovaSeq 6000 platform using for the read 1 28 cycles, i7 index 10 cycles, i5 index 10 cycles and read 2 90 cycles.

### Acute brain slice preparation

Mice of postnatal day 13–15, 20-23 or 37-40 were anesthetized with isoflurane, decapitated, and brains were removed quickly and submerged into ice-cold artificial CSF (aCSF) (composition in mM for mice up to P23: 118 NaCl, 25 NaHCO_3_, 3 KCl, 1 NaH_2_PO_4_, 1.5 CaCl_2_, 1 MgCl_2_, 30 glucose, composition in mM for mice P37-P40: 118 NaCl, 25 NaHCO_3_, 3 KCl, 1 NaH_2_PO_4_, 1 CaCl_2_, 1 MgCl_2_, 1 glucose, 29 sucrose) saturated with 95% O_2_, 5% CO_2_, at pH 7.4. After cutting the midline, 350 µm thick coronal sections containing the dorsal hippocampus for whole cell recordings were obtained using a vibratome (Microm, H650 V, Leica Microsystems). For two-photon calcium imaging, 350 µm thick horizontal sections were cut instead. After slicing, the brain slices were recovered in recording aCSF at 36°C in slicing aCSF for 45 min and afterwards stored at room temperature in recording aCSF. For recording, acute slices were placed in an upright microscope (Olympus, Japan) equipped with immersion differential interference contrast objectives (5x, 60x) coupled to a monochromatic camera (XM10, Olympus, Japan) and maintained at a temperature of 33 ± 1°C while perfusing with oxygenated recording aCSF.

### Electrophysiological recordings

Experiments were performed in current-clamp and in voltage clamp mode using an Axopatch 200B amplifier connected to a Digidata 1440A (both Molecular devices, USA). In current-clamp active and passive membrane properties of patched neurons were recorded using electrodes with resistances ranging from 3-6 MΩ with the following intracellular solution (in mM): 140 K-gluconate, 1 CaCl_2_, 10 EGTA, 2 MgCl_2_, 4 Na_2_-ATP, 10 HEPES, pH 7.2; the osmolarity was 290-300 mOsmol. Biocytin was added to the intracellular solutions for current-clamp recordings at a concentration of 0.5% (w/v) to confirm cell type post-hoc by Streptavidin counterstaining. To evoke trains of action potentials and measure sag-potentials, neurons were clamped to - 55mV, effectively −70 mV, taking a liquid junction potential of + 15 mV into account. Spontaneous and optically stimulated inhibitory synaptic currents were recorded in presence of 50µM CNQX and 25 µM AP5 (with additional 1 µM TTX to record miniature inhibitory currents) using a patch pipette filled with 105 CsCl, 35 CsOH x H_2_O, 10 HEPES, 10 EGTA, 10 Phosphocreatine, 4 Mg-ATP, 0.3 Na-GTP, 14 D-Mannitol pH 7.3, with CsOH; the osmolarity was 290-300 mOsmol, leading to an increase in driving force of chloride ions and therefore reversing IPSC polarity at the holding potential of −70 mV. The currents were filtered online at 10 kHz and recorded with a sampling rate of 20 kHz for 120 seconds. Cell capacitance and series were compensated, the series resistance was corrected by 85%. Neurons with a series resistance > 25 MΩ were discarded from analysis. During voltage-clamp recordings, cells were clamped to −70 mV. For stimulation of oIPSCs a Polygon 1000 pattern illuminator (Mightex, USA) coupled to a 470 nm Super High-Power LED Collimator as light source (6300 mW^2^ maximal output power, Mightex, USA) was used for illumination. Illumination patterns at 50% total LED power with pulse length of 5 ms with 5 separate pulses at pulse frequencies of 10 or 40 Hz were designed in Polyscan2 and triggered via a TTL signal by the digitizer. For somatic stimulation, mCherry-positive somata were illuminated, while perisomatic stimulation was performed by illumination of the patched postsynaptic PC’s soma.

### Immunohistochemistry (IHC) and confocal and Imaging

For neuronal morphology reconstruction or IHC staining with non-synaptic markers, brain slices were fixed after recordings overnight in 4% paraformaldehyde (Morphisto GmbH, Offenbach am Main, Germany), followed by washing in DPBS (Thermo Fisher Scientific, Waltham, MA, USA, Cat. 12559069) and blocking with 1% normal goat serum and 0.2% Triton X-100 for 2 hours. Subsequent primary antibody incubation was performed overnight at 4°C with the following antibodies and dilutions: Chicken anti-mCherry 1:250, Aves labs; mouse anti-parvalbumin 1:1000, Sigma. For visualization of filled cells, counterstaining with Streptavidin-Cy3, 1:100, Sigma, Cat. S6402 for three hours at room temperature was performed instead. For staining of synaptic proteins 150 µm thick fresh brain slices were fixed in 0.5% methanol and 1% PFA in PEM buffer for one hour at 4°C instead. Slices were subsequently incubated in 0.1 M citrate buffer PH 4.5 overnight and antibody binding epitopes were recovered by microwaving slices submerged in citrate buffer for 30 seconds. After blocking with 1% normal goat serum and 0.2% Triton X-100 for two hours at room temperature staining with following primary antibodies was performed: rabbit anti-ankyrinG 1:1000, Synaptic systems; mouse-anti Gephyrin 1:500, Synaptic systems, clone 3B11; chicken-anti VGAT 1:500, Synaptic systems; mouse anti-synaptotagmin 2 1:1000, Oregon Zebrafish facility. After washing in PBS 3 x 15 minutes, sections were counterstained with the following secondary antibodies: goat anti-rabbit IgG Alexa Fluor-488, 1:200; goat anti-chicken IgG Alexa Fluor-568, 1:200; goat anti-mouse IgG Alexa Fluor-647, 1:200. After three additional washing steps, slices were mounted with DAPI Fluoromount-G (Southern Biotech, Birmingham, AL, USA, Cat. 011-20). Immunohistological confocal images were acquired on an upright Leica SP8 microscope (Leica, Germany).

### Two-photon calcium imaging of brain slices

Calcium imaging from acute brain slices was performed using an upright Trimscope II (Miltenyi Biotec) equipped with a resonant scanner and a Leica HC FLUOTAR L 25x/1,0 IMM (ne=1,457) objective. Gcamp7f was excited with an Alcor 920 nm wavelength-locked femtosecond-pulsed laser with an average output power of 1,7 watts at 80 MHz (Spark Lasers). Images were acquired at 30.5 Hz using a beam power of 40-60%. Emitted fluorescence was recorded on a high sensitivity photomultiplier module (Hamamatsu) with a gain of 80% after passing through a 490/20 nm band pass filter. During recordings, horizontal brain slices were submerged in recording aCSF (118 NaCl, 25 NaHCO_3_, 8 KCl, 1 NaH_2_PO_4_, 1.5 CaCl_2_, 0.5 MgCl_2_, 30 glucose, saturated with 95% O_2_, 5% CO_2_, at pH 7.4) and temperature was maintained at 34°C. Ca-imaging movies were recorded for three minutes 15 minutes after placing the brain slice in the microscope setup.

### Transmission electron microscopy

For transmission electron microscopy, mice were anesthetized with a mixture of ketamine (120 mg/kg) and xylazine (8 mg/kg) and transcardially perfused with 4% PFA/4% glutaraldehyde (GA) in PBS. Neocortex and hippocampus was post-fixed in the same fixative for 4 h. After washes in 0.1 M cacodylate buffer (pH 7.4), samples were post-fixed and contrast was added by using 1% OsO_4_, dehydrated in an ethanol series (50, 70, 96, and 100%) during which the 70% ethanol was saturated with uranyl acetate for contrast enhancement. After the dehydration was completed by incubation in propylene oxide, specimens were embedded in Araldite (Serva, Heidelberg, Germany) which was hardened at 60 °C for 48 h. 400 nm semi-thin sections and 50-60 nm ultra-thin sections were cut on an FCR Reichert Ultracut ultramicrotome (Leica, Bensheim, Germany). The semi-thin sections were dried on glass slides, and sections were stained for 1 min with Richardson’s solution at 70 °C and rinsed with distilled water. From the selected region, ultra-thin sections were made, mounted on Pioloform-coated copper grids, and contrasted with lead citrate. Images were acquired with an EM10A electron microscope (Carl Zeiss, Oberkochen, Germany) and a digital camera (Tröndle, Germany). No further image modification was applied.

## Data analysis

### Electrophysiology

Optically stimulated synaptic events, action potential based analysis, passive cellular properties and τ_decay_ of synaptic events were analyzed using Clampfit 11.2 (Molecular devices). Spontaneous synaptic events were detected automatically after baseline correction by a self-written script in Spike2 version 10 (CED) taking a minimum event amplitude of 20 pA as cutoff as well as a minimum acceleration of −30000 pA/s^2^ within a 2 ms time window and a minimum lag time between individual events of 5 ms. Events with amplitudes larger than 2000 pA were regarded as non-synaptic and discarded from analysis. τ_decay_ was calculated by fitting the falling slope of the synaptic event with a single exponential function. For cumulative distribution plots all synaptic events recorded from one cell were used to obtain cell-specific cumulative distributions and then averaged with the data from all other cells of the same genotype. The resting membrane potential (V_m_) was evaluated in current-clamp mode two minutes after rupturing the patch without applying any current. Input resistance (R_IN_) was calculated by plotting a linear regression curve over the membrane voltage obtained after applying hyperpolarizing current steps ranging from −10 to −110 pA with the slope of the curve reflecting R_IN_. For AP analysis in trains, excitatory PCs and regular-spiking interneurons were stimulated with 800 ms long step current injections ranging from −200 to 300 pA while fsIN were stimulated with 800 ms long step current injections from −200 to 800 pA. For PCs, firing frequencies were compared only for the first 150 ms of current injection due to occurrence of plateauing at higher current injections. AP parameters for rsIN and fsIN were analyzed for the last AP in the train at 150 pA current injection after the neuron had reached steady state firing while for PCs the first AP was analyzed due to plateauing of the membrane potential at the same current injection step. AP threshold was determined as the membrane potential recorded when the first derivative reached a speed of 20 mV/s during the rising phase of the AP. AP amplitude and rise time were measured as voltage change/time interval from threshold to the peak, while the repolarisation amplitude was calculated as voltage change/time interval from AP peak to anti-peak during the falling phase. AP halfwidth was defined as the time interval between the points when the AP waveform reached the half-maximal amplitude during the rising and falling phase of the AP. AP rheobase was defined as the minimal current step needed to elicit at least one AP. Sag potential elicited at −300 pA current injection was determined as the mean change of voltage between the initial and the final voltage during current injection. Phase plots were generated from the last AP in train at 150 pA current injection for fsIN, starting 0.2 ms prior to AP threshold for a total time window of 1.5 ms with 0.1 ms bins.

### Ca-Imaging

Calcium-imaging data acquired by two-photon microscopy was analyzed with the Mesmerize 0.9.0 software pipeline (Kolar et al. 2021): Movies were first motion-corrected using the non-rigid motion correction implementation, followed by ROI detection via CNMF both from the CaImAn library (Giovannucci et al. 2019) with estimated decay-time for Gcamp7f signals of 0.4 seconds for each calcium transient. After ROI detection, fluorescence signals were converted to dF/F_0_ signals and single spike data were extracted by extrapolation. For calcium-peak detection, dF/F_0_ signals were first filtered by a Butterworth filter of second order and a normalized cutoff frequency of 15.25 Hz. The original dF/F_0_ signal, the filtered signal, a filtered signal normalized to the maximum peak and its derivative were used for peak detection with inbuilt PeakDetect function. For peak parameters, calcium-peak duration was calculated as the time between the left and right base flanking the peak while for peak AUC the integral was determined as a second parameter to assess neuron burst behavior. Peak amplitude was calculated as the amplitude difference between the left base and the signal peak. Peak frequency was calculated as the mean of instantaneous frequency of peaks in one signal trace.

### Analysis of EM pictures

EM data were analyzed within Fiji (Schindelin et al. 2012). Hereby, the synaptic length was measured tracing the postsynaptic density while the synaptic width was measured from the edge of the membrane density from pre-to postsynaptic side.

### Single nuclei RNA sequencing analysis

Samples were demultiplexed using Illumina’s bcl2fastq conversion tool and the 10x Genomics pipeline Cell Ranger count (v6.0.1) to perform alignment against the 10x Genomics pre-built Cell Ranger reference GRCh38-2020-A (introns included), filtering, barcode counting, and UMI counting. Additionally, Chromium cellular barcodes were used to generate (filtered) gene-barcode matrices, and later determine clusters, and perform gene expression analysis. Gene barcode matrices from every mouse sample were imported to RStudio and separately pre-processed using Seurat 4.3.0 (Hao et al. 2021) as follows: Counts of genes appearing in less than three nuclei and nuclei with less than 1000 or more than 7500 unique genes, a higher percentage of 15% mitochondrial counts, or a total number of molecules lower than 1000 per nuclei were discarded from analysis. Counts from remaining nuclei were normalized, scaled, and analyzed via PCA based on the top 500 highly variable genes, identified beforehand via variance-stabilizing transformation method. The top 15 PCAs explaining the largest standard deviation between cells were used to simulate a dataset of nuclei doublets using package DoubletFinder 2.0.3 (McGinnis et al. 2019) and nuclei doublets in the real dataset were removed expecting a doublet percentage ratio of 8%. After doublet elimination data from all six animals were combined into one Seurat object and integrated as a separate assay-object based on the samples genotype using the inbuilt Seurat RPCA integration anchor algorithm. This later allowed clustering of identical cell types together despite genotype dependent changes in gene expression. To obtain optimized cell cluster resolution representing the different cell types, top 15 PCA dimensions of integrated nuclei datasets were clustered with the findNeighbour function and the resolution package Clustree 0.5.0 (Zappia und Oshlack 2018) before dimensionality reduction via UMAP. Cell types were subsequently annotated by hand based on literature established hippocampal marker genes, especially the Allen brain map transcriptomics explorer using the “Mouse – Whole Cortex & Hippocampus - 10x” dataset (Yao et al. 2021).

To test for differentially expressed genes (DEG) between genotypes, Wilcoxon ranking test was used on each identified cell cluster while DeSeq2 test was used to generate DEG lists to perform further GSEA analysis based on the package Fgsea 1.18.0 (Korotkevich et al. 2016) for all three classes of gene ontology (GO) terms using the GO annotation list “m5.go.v2023.1.Mm.symbols.gmt” from gsea-msigdb.org. In contrast to classical GO-term annotation, GSEA takes into account not only the GO annotation of each gene, but also the log2 fold change and the polarity of gene dysregulation. For enrichment analysis, gene sets consisting of less than 15 genes, or more than 400 genes were omitted before collapsing enriched gene sets to pathways. Collapsed pathways with a p value cutoff of 0.05 for enrichment were then considered as significantly altered between genotypes.

Pseudotime analysis of selected cell clusters was performed using the package Monocle3 1.3.1 (Trapnell et al. 2014; Qiu et al. 2017): Nearest neighbor properties of all nuclei of chosen clusters were first re-analyzed by on PCAs including either all highly-variable genes or a subset of genes (see Results section). Created UMAP plots of resulting dimensionality reduction were then assigned to one partition and pseudotime trajectories were learned by the system without additional rounds of pruning, creating additional branches and retaining branches otherwise removed. Start points with pseudotime 0 were chosen manually on the created trajectory based on either branch ends with the highest ratio of data points from the P16 dataset (interneuron pseudotime analysis) or the highest ratio of wt nuclei (pyramidal cell P20 pseudotime analysis). Testing for differentially expressed genes along the created trajectories was done with Moran’s I test; genes with q < 0.05 and Morans’s I > 0.1 were regarded as genes with significant influence on the created trajectories.

To analyze underlying changes in TF activity in selected cell clusters, we performed gene regulatory network (GRN) analysis on selected neuron populations with the SCENIC 1.3.1 package (Aibar et al. 2017). Before GRN building, genes were filtered to be expressed in at least 0,1% of cells and to have a minimum summed up gene count larger than 1% of the total cell number. Gene expression was mapped onto the RcisTarget motif databases mm9-tss-centered-10kb-7species.mc9nr.feather and mm9-500bp-upstream-7species.mc9nr.feather. Two separate GRNs for inhibitory and excitatory cell clusters in the P20 dataset were built using algorithm GRNBoost2 (Moerman et al. 2019) on the log2 transformed gene expression matrix. To identify differentially regulated regulon activity between genotypes, the GRN was mapped back onto the original Seurat object as a separate assay object and Wilcoxon ranking test was used on AUC for each regulon activity for each cell cluster spit by genotype.

### Statistical analysis

Statistical analysis was conducted in Graphpad Prism 10. All values displayed in text and graphs of this manuscript are mean ± standard error of the mean (SEM if not otherwise stated. All statistical significance was tested by standard Student’s t-test for normally distributed data or Mann-Whitney test for non-normally distributed data. Significant differences are marked as follows: *p ≤ 0.05, **p ≤ 0.01, ***p ≤ 0.001. All data analysis besides analyses of RNA sequencing were performed by researchers blinded to the genotype. Detailed information about statistical values can be found in the supplementary tables.

## Supporting information

Supplementary statistic table

## Acknowledgements

We thank Gabriele Frommer-Kästle for the technical assistance with EM sample preparation. We thank the NGS Competence Center Tübingen especially Dr. Nicolas Casadei for organizing and conducting snRNA sequencing and data preparation for downstream analysis. In addition, we thank Dr. Malte Stockebrand for sharing and optimizing the nuclei isolation protocol and further members of the FOR2715 unit for advice on RNA-sequencing downstream analysis. We thank Ana Fulgencio Maisch for the technical assistance with animal breeding and genotyping. Graphical figure elements were created with Biorender.com.

## Author contribution

NL and PM performed sample preparation and data analysis for snRNA seq data. NL, MS, and MA performed patch-clamp experiments and IHC. NL performed two-photon Ca-imaging experiments. FP, PFB, AS and NL performed sample preparation for electron microscopy. FK and NL implemented coding for Ca-imaging data analysis. NL, TVW, PM and SI wrote the manuscript. HK, UBSH and HL helped with data interpretation and manuscript preparation. NL, TVW, PM and HL designed the study. TVW and NL developed *in vivo* methodology. Project administration, TVW. All authors contributed to the article and approved the submitted version.

---

**Supplementary Figure 1:**
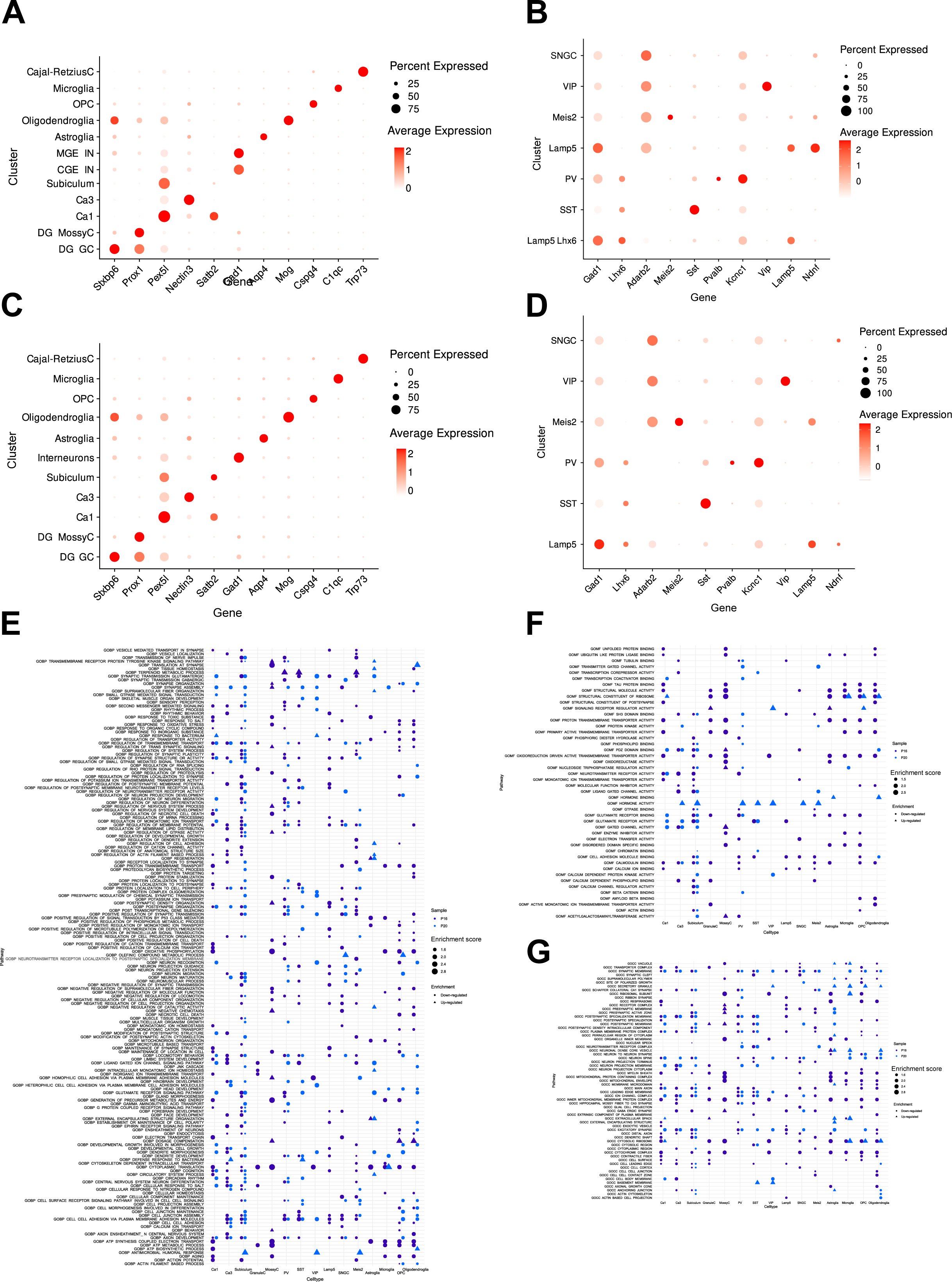
Extended RNA seq analysis of hippocampal cell populations over pathodevelopment. **A-D**: Expression of marker genes in identified cell clusters as dot plots at P16 (A, B) and P20 (C, D). Interneuron subtypes (B, D) could only be identified after additional sub-clustering of GABAergic neuron clusters. **E-G**: Complete list of enriched GO-terms after GSEA analysis in neuronal and glial cell populations at P16 and P20. The lists are divided by the three main GO terminology branches “Biological process” (E), “Molecular function” (F) and “Cellular compartment” (G).

**Supplementary Figure 2:**
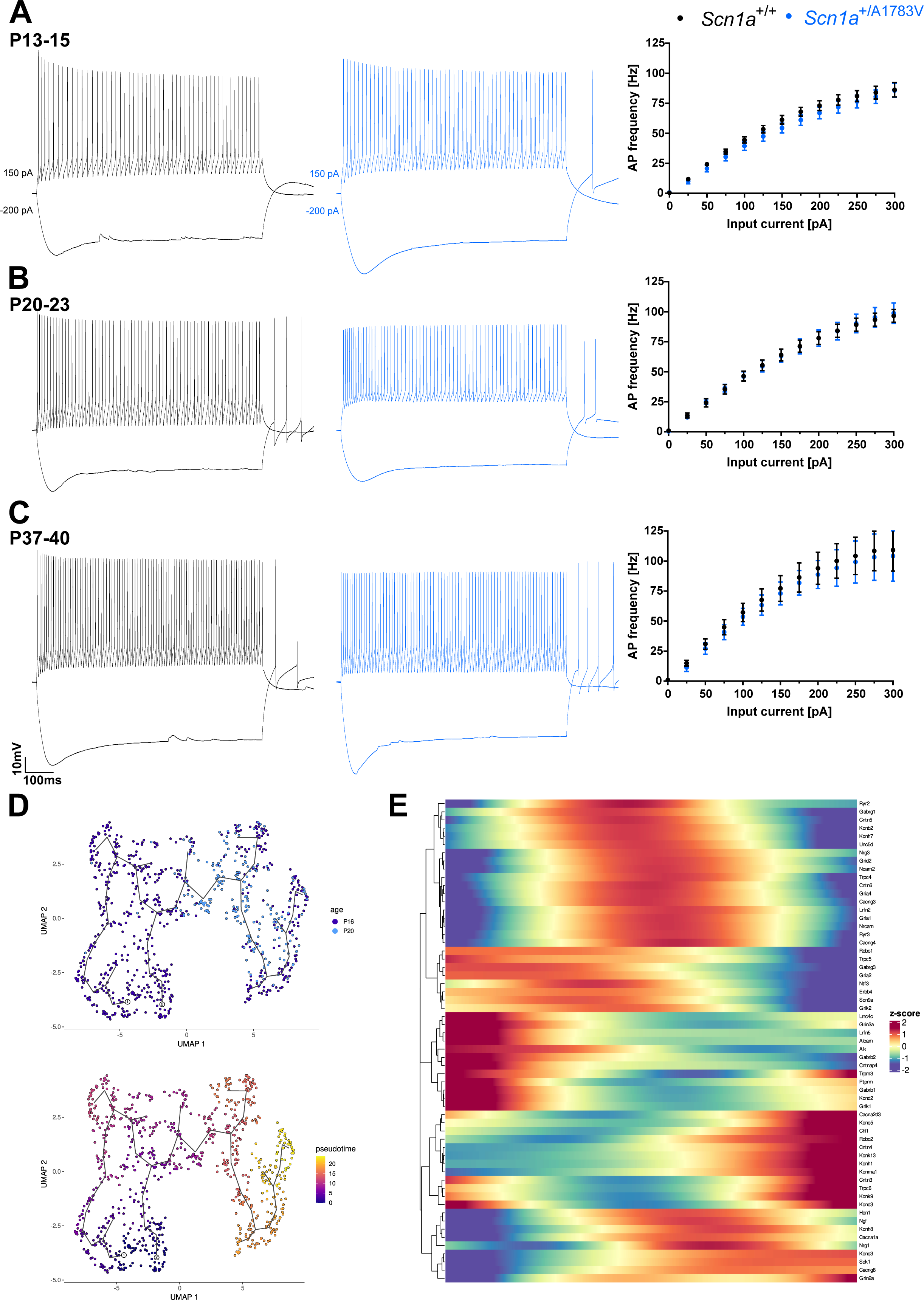
CA1 regular-spiking-interneuron excitability over pathodevelopment. **A-C:** Representative AP trains at −200 pA and 150 pA current injection over a duration of 800 ms and frequency-current curves of for rsINs of wt (black) and *Scn1a*^+/A1783V^ (blue) mice before seizure onset, at the high-seizure phase and the stabilization disease phase. There was no change in somatic excitability between wt and *Scn1a*^+/A1783V^ cells. **D**: Umap representations of hippocampal SST interneurons of *Scn1a*^+/A1783V^ mice at P16 and 20 sub-clustered based on all dysregulated genes at P20 labeled as function of pseudotime and age using Monocle 3. **E**: Heatmap of genes describing the SST cell pseudotime trajectory with a cutoff of Moran’s i > 0.1 and a q-value > 0.05 (x-axis indicates pseudotime). Genes were clustered based on the pattern of change in expression along the pseudotime trajectory. Positive z-Scores display peak expression of listed genes along pseudotime while negative z-Scores reflect downregulation of particular genes along pseudotime.

**Supplementary Figure 3:**
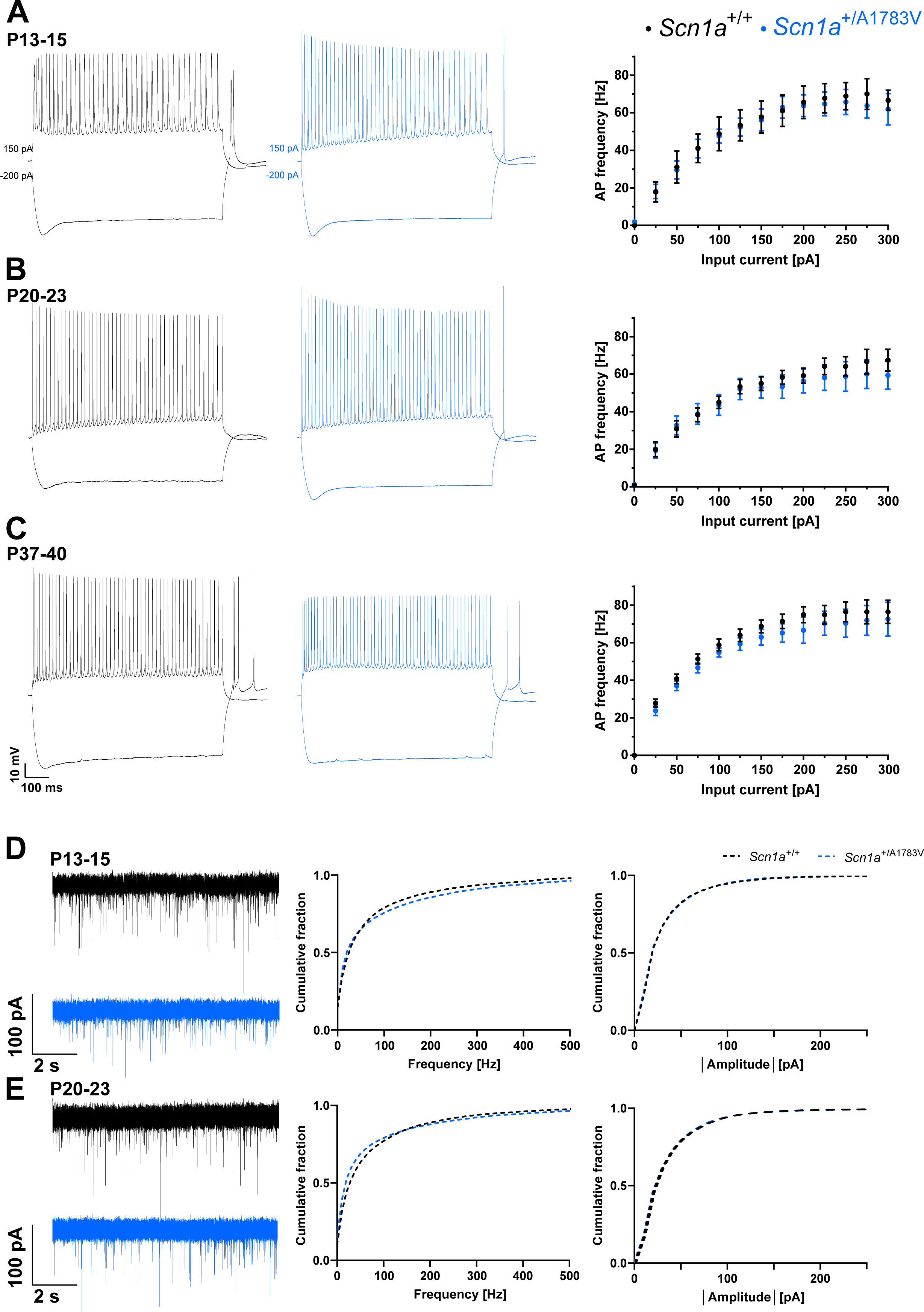
Changes of CA1 pyramidal excitability over pathodevelopment. **A-C:** Representative AP trains at −200 pA and 150 pA current injection over a duration of 800 ms and frequency-current curves of CA1 PCs of wt (black) and *Scn1a*^+/A1783V^ (blue) mice before seizure onset, at high-seizure and stabilization disease phase. There was no change in somatic excitability between wt and *Scn1a*^+/A1783V^ cells. E: Representative AP waveforms of fsIN at the start (left panel) and end (right) of an 800 ms long current injection of 150 pA in wt and *Scn1a*^+/A1783V^ mice before seizure onset, during high frequency-seizure and stabilization phase. **D, E**: Representative traces of spontaneous excitatory postsynaptic currents (sEPSC)s recorded from fsINs in CA1 at P13-15 and P20-23 before seizure onset in *Scn1a*^+/+^ and *Scn1a*^+/A1783V^ mice. **B-D**: Cumulative fraction plots of event instantaneous frequency and amplitude. SEPSC frequency and amplitudes were not altered over phenotypic disease onset in *Scn1a*^+/A1783V^ mice.

**Supplementary Figure 4:**
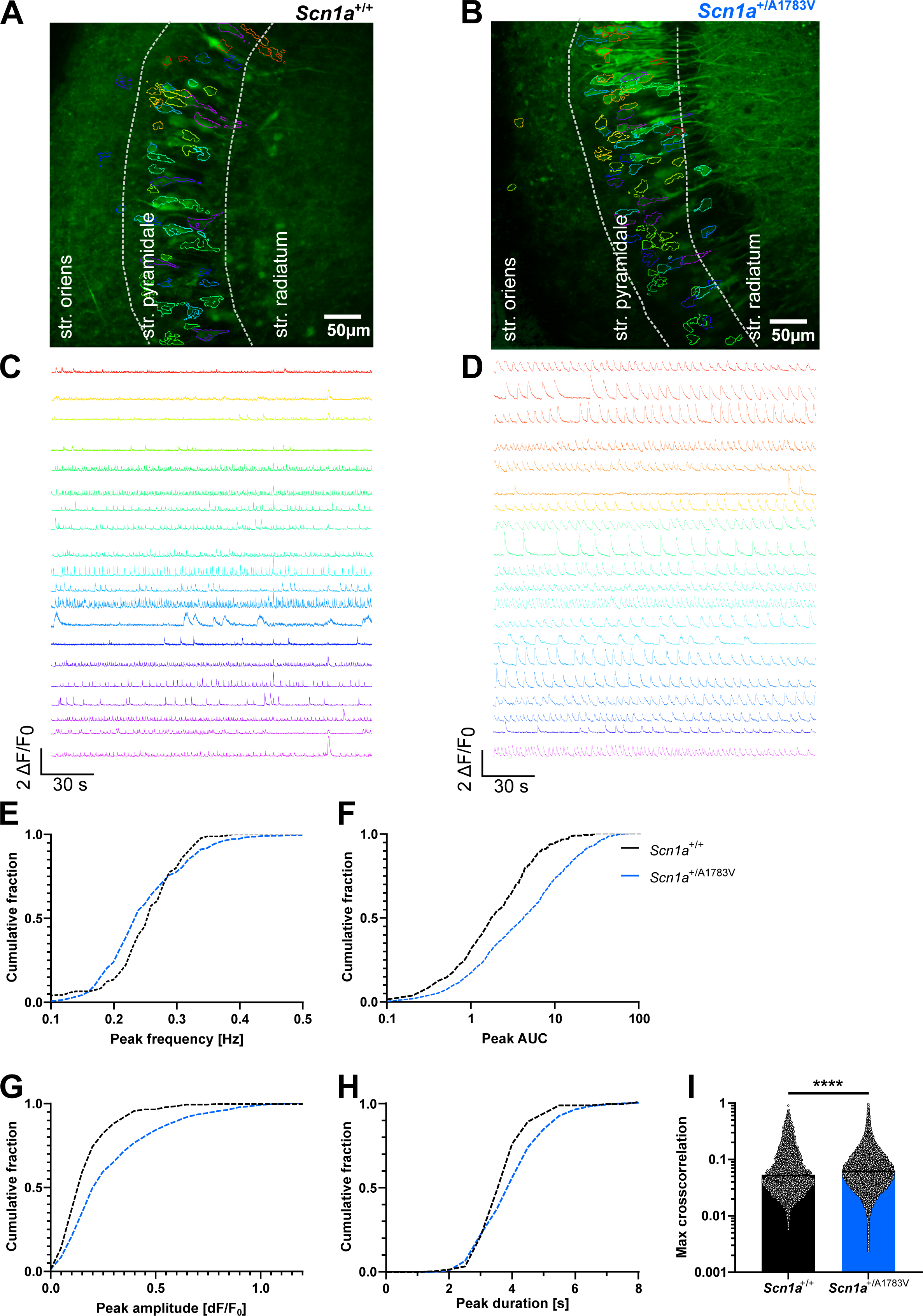
Hyperexcitability of CA3 neurons revealed by two-photon Ca-imaging in horizontal hippocampal brain slices. **A, B**: Representative field of view of a two-photon calcium imaging movie collapsed by orthogonal projection of movie standard deviation. Colored regions of interest (ROIs) in the CA3 *str. pyramidale* of *Scn1a*^+/+^ and *Scn1a*^+/A1783V^ mice at P20-23 detected by the CNMF algorithm coincide with the colored traces in C and D. **C, D**: Extracted calcium signals of GCaMP7f calcium sensor as ΔF/F signals from movie recordings over a period of three minutes in 0.5 mM Mg^2+^ and 8 mM K^+^ aCSF. **E-H**: Cumulative fraction plots of Ca-peak frequency, AUC, peak amplitude, and duration, comparing calcium-traces wt and *Scn1a*^+/A1783V^ cells. *Scn1a*^+/A1783V^ cells displayed a significant lower event frequency, while event AUC, amplitude and duration were significantly enlarged. **I**: Bar graph of maximal cross-correlation between neuronal calcium-traces in a 500 ms lag window in *wt* and *Scn1a*^+/A1783V^ slices. Bar represents the median. Calcium signals were significantly more synchronous between single neurons in brain slices from *Scn1a*^+/A1783V^ mice.

**Supplementary Figure 5:**
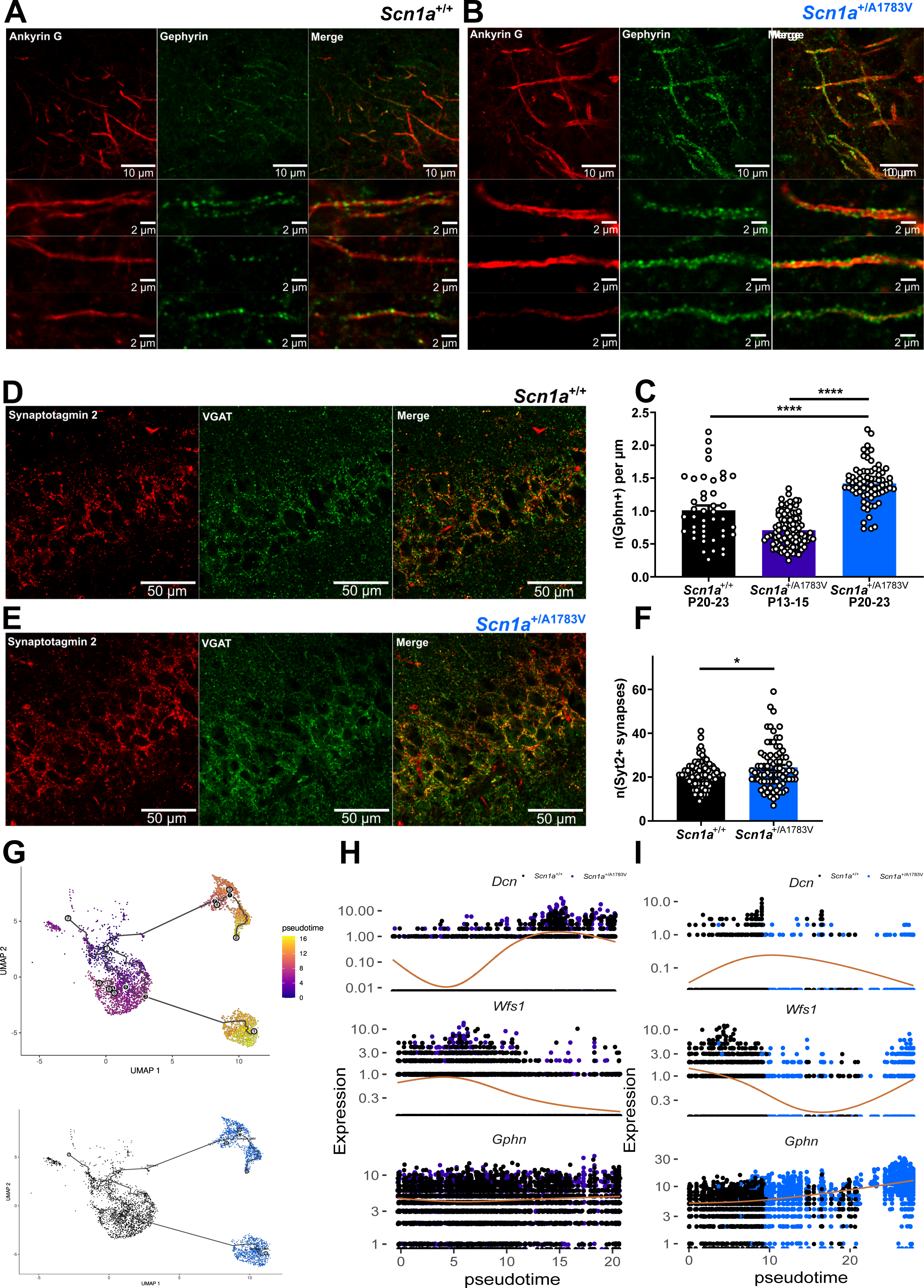
Immunohistological staining and quantification of GABAergic synapses in CA1 *str. pyramidale*. **A, B**: Representative immunohistological staining of AIS marker ankyrinG (red) and inhibitory postsynaptic density marker gephyrin (green) in *Scn1a*^+/+^ and *Scn1a*^+/A1783VV^ mice at P20-23 in CA1. Gephyrin puncta colocalize with the axon initial segment of PCs. **C:** Quantification of gephyrin puncta normalized to AIS length. AIS of *Scn1a*^+/A1783V^ mice at P20-23 displayed a significantly increased density of inhibitory synapses compared to *Scn1a*^+/+^ mice at P20-23 and *Scn1a*^+/A1783V^ mice at P13-15. **D, E**: Representative immunohistological stainings of inhibitory presynaptic vesicular GABA transporter (VGAT, green) and PVIN presynaptic terminal marker protein synaptotagmin 2 (Syt2, red) in *Scn1a*^+/+^ and *Scn1a*^+/A1783V^ mice at P20-23 in CA1. Syt2 puncta colocalize with VGAT puncta at perisomatic inhibitory synapses onto PCs in CA1. **F:** Quantification of Syt2 puncta per soma. Somata of PCs in *Scn1a*^+/A1783VV^ mice at P20-23 displayed a significantly increased density of Syt2^+^ synapses compared to *Scn1a*^+/+^ mice. **G**: Umap representations of CA1 PCs subclustered at P16 and P20 labeled as function of pseudotime and cell genotype using Monocle 3. **H, I**: Expression of ventral CA1 marker *Dcn*, dorsal CA1 marker *Wfs1* and inhibitory synapse marker gephyrin over pseudotime and labeled by genotype in CA1 PCs at P16 (H) and P20 (I). There was an upregulation of *Gephn* expression at P20 in dorsal CA1 PCs in *Scn1a*^+/A1783V^ mice while there was no change in *Gephn* expression before seizure onset in dorsal and ventral CA1 PCs.

